# Proliferation and differentiation of intestinal stem cells depends on the zinc finger transcription factor BCL11/Chronophage

**DOI:** 10.1101/2024.09.08.611891

**Authors:** Siamak Redhai, Nick Hirschmüller, Tianyu Wang, Shivohum Bahuguna, Svenja Leible, Stefan Peidli, Erica Valentani, Sviatoslav Kharuk, Michaela Holzem, Lea Bräckow, Fillip Port, David Ibberson, Wolfgang Huber, Michael Boutros

**Author notes:** Contributed equally (Siamak Redhai, Nick Hirschmüller). Cardiovascular Epidemiology Unit, Department of Public Health and Primary Care, University of Cambridge, Cambridge, UK.

## Abstract

The molecular programs that drive proliferation and differentiation of intestinal stem cells (ISCs) are essential for organismal fitness. Notch signalling regulates the binary fate decision of ISCs, favouring enterocyte commitment when Notch activity is high and enteroendocrine cell (EE) fate when activity is low. However, the gene regulatory mechanisms that underlie this process on an organ scale remain poorly understood. Here, we find that the expression of the C2H2-type zinc-finger transcription factor *Chronophage* (*Cph*), homologous to mammalian BCL11, increases specifically along the ISC-to-EE lineage when Notch is inactivated. We show that the expression of *Cph* is regulated by the Achaete-Scute Complex (AS-C) gene, *scute,* which directly binds to multiple sites within the *Cph* locus to promote its expression. Our genetic and single-cell RNA sequencing experiments demonstrate that Cph maintains the ISC and EE populations and is necessary to remodel the transcriptome of progenitor cells with low Notch activity. By identifying and functionally validating Cph target genes, we uncover a novel role for *sugar free frosting* (*sff*) in directing proliferative and lineage commitment steps of ISCs. Our results shed light on the mechanisms by which *Cph* sustains intestinal epithelial homeostasis and could represent a conserved strategy for balancing proliferation and differentiation in different tissues and species.

## INTRODUCTION

Replenishment of epithelial cells in adult tissue is central for homeostatic balance^1^. The intestinal epithelium consists of diverse cell types which are derived from multipotent intestinal stem cells (ISCs). The process of differentiating into lineage-specific cell types depends on multiple factors including asymmetric cell division, detachment from the ISC niche, mechanical regulation and signalling dynamics^2–4^. Such mechanisms are utilised to ensure that turnover of epithelial cells is faithfully coordinated during environmental stress, tissue damage and to maintain homeostasis. The majority of the intestinal epithelium consists of enterocytes (ECs) which are primarily responsible for absorptive functions^5^. This is in contrast to the neuropeptide secreting enteroendocrine cells (EEs), which comprise a small population of epithelial cells that regulate endocrine processes, including food intake, appetite, gut motility and metabolism^6,7^.

The *Drosophila melanogaster* midgut has proven to be an excellent model to understand ISC fate determination, with many of the biological mechanisms conserved in mammals^8–10^. The fly midgut consists of self-renewing multipotent ISCs that are scattered along the intestine and give rise to Enteroblast (EBs) and Enteroendocrine progenitors (EEPs) which terminally differentiate into ECs and EEs, respectively^11–13^. Recent studies have identified 10 subpopulations of enteroendocrine cells (EEs) that are spatially confined to specific regions of the midgut^14–16^. These subpopulations are broadly categorised into three major groups according to the expression of distinct neuropeptides; class I: AstC^+^ cells, class II: Tk^+^ cells, and class III: CCHa2^+^ cells. Notch signalling plays an important role in determining the fate of ISCs, favouring ECs when Notch activity is high, and EEs when Notch activity is low^17–19^. Inactivation of the Notch pathway in the *Drosophila* intestine results in excessive proliferation and accelerates the formation of AstC^+^ EEs, leading to tumour-like structures that develop mainly in the posterior midgut, and to a decline in survival^20,21^.

Previous studies highlighted a number of transcription factors (TFs) that regulate commitment to an EE fate. For example, Notch signalling negatively regulates the expression of the Achaete-Scute Complex (AS-C) basic helix–loop–helix TF *scute* (*sc*)^22,23^. When Notch signalling is low, *sc* is activated through a transcriptional self-stimulatory loop which results in asymmetric cell division and terminal differentiation of ISCs into EEs^22^, a process that is regulated by the homeobox TF *prospero* (*pros*), which governs the differentiation program in EEs^15^. Upstream TFs such as *klumpfuss* (*klu*) normally suppress *sc* expression and regulate apoptosis to drive EC lineage commitment^24,25^. Similarly, the TF *tramtrack* (*ttk*) also functions as a negative regulator of *sc* expression^26^. The Ttk protein is kept in check by the E3 ubiquitin ligase Seven in absentia (Sina) and the adaptor protein Phyllopod (Phyl) which work together to ubiquitinate Ttk, leading to its degradation by the proteasome^27^. Despite these findings, our understanding of the regulatory networks determining the EEs lineage on an organ scale remain poorly understood.

In this study we profiled the transcriptome of 46,799 single cells from the adult *Drosophila* midgut during homeostasis and multiple perturbed conditions involving *Notch* and *Cph*. We identified the C2H2 zinc-finger TF, *Chronophage* (*Cph*) as a crucial regulator of progenitor cell maintenance and EE fate commitment. *Cph* expression is induced early during the ISC-to-EE lineage when Notch activity is diminished and is required for generating class I and II EEs. Chromatin binding profiles show that Sc binds directly to the *Cph* locus and promotes its expression. scRNA-seq of single and double perturbation experiments between *Notch* and *Cph* demonstrate that *Cph* is required for transcriptional reprogramming of ISCs and EEPs when Notch is depleted. By profiling Cph chromatin binding sites, we identified a number of key target genes, including *sff* which directs proliferative and lineage commitment steps of ISCs. Our findings therefore highlight a previously uncharacterised gene regulatory network centred on *Cph* that tunes intestinal epithelial identity.

## RESULTS

### scRNA-seq of *Notch* mutant intestinal cells reveals major changes in cell type composition

To explore how different intestinal cell types respond to impairment of Notch signalling, we performed single-cell RNA sequencing (scRNA-seq) of the adult *Drosophila* intestine at homeostatic condition and under progenitor-specific CRISPR mutagenesis of the Notch receptor with two single guide RNA (*sgRNA^x^*^2^) (herein referred to as *Notch^sgRNAx^*^2^)^28^ using the *esg^TS^* driver system (*esg-Gal4, tub-Gal80^TS^>GFP, Cas9^p.^*^2^) (Fig. 1a). The *esg^TS^* driver permits temporally limited expression of transgenes and GFP specifically within the progenitor population of the intestine. With each condition assayed in duplicate, we obtained a total of 29,741 single cell gene expression profile with high data quality (control: 13,452 cells; *Notch^sgRNAx^*^2^: 16,289 cells), surpassing previously published scRNA-seq datasets for the *Drosophila* intestine^14,29^ (Extended Data Fig. 1a-d). As a result, we provide to the community a Shiny app to enable visualisation and exploration of our scRNA-seq dataset: https://shiny-portal.embl.de/shinyapps/app/16_IntestiMap

**Fig. 1.**
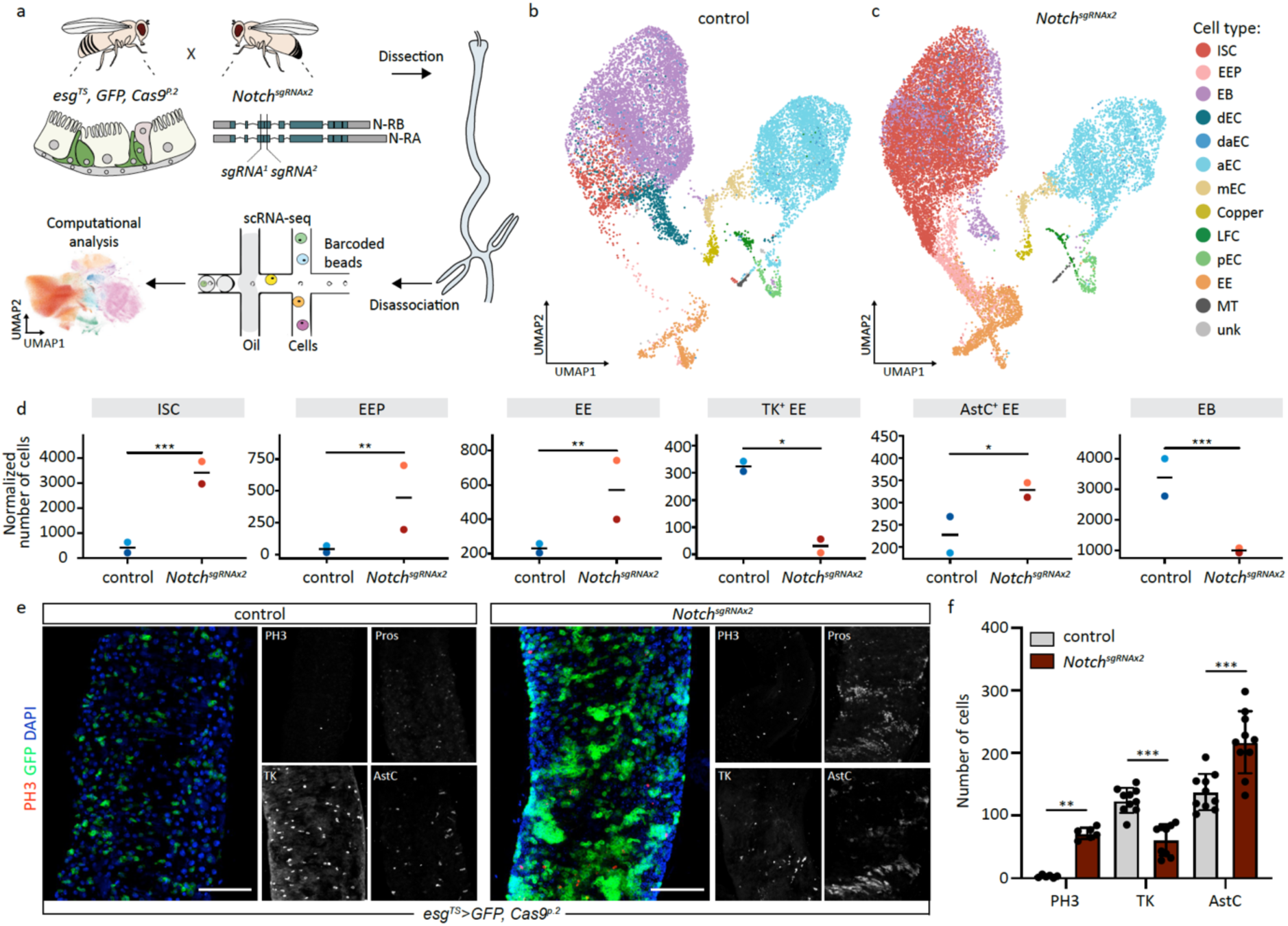
scRNA-seq of intestinal cells reveals major changes in cell type composition upon *Notch* inactivation. **(a)** Schematic of single cell RNA-sequencing experimental workflow. CRISPR mutagenesis of *Notch* was achieved by expressing two *sgRNA* targeting the *Notch* locus with the *esg^TS^, GFP, Cas9^p.2^* system, which drives expression of *sgRNA*, *Cas9^p.2^* and *GFP* within the progenitor population. **(b, c)** Uniform Manifold Approximation and Projection (UMAP) plot of combined scRNA-seq dataset for control condition **(b)** and Notch perturbed condition **(c)**. Clusters were manually annotated with guidance from previously published scRNA-seq datasets for the *Drosophila* intestine. **(d)** Quantification of cell type abundance coloured according to scRNA-seq replicates. **(e)** *in vivo* CIRSPR mutagenesis of *Notch* in progenitor cells results in an expansion of the progenitor population, an increase in the number of PH3^+^ mitotically active cells, an increase in AstC^+^ EEs and a decrease in TK^+^ EEs. **(f)** Quantification of different cell types in control and *Notch* mutant conditions. One-way ANOVA test with Tukey post hoc comparison was used for (F). \**P* < 0.05, \*\**P* < 0.01, \*\*\**P* < 0.001. Scale bar for e is 100 μm.

We clustered the cells’ transcriptome profiles, taking guidance from previous cell type catalogues^14,29^ and manually categorised our data into 10 major cell types, including ISCs, EBs, EEPs, ECs, and EEs and their respective subtypes (Fig. 1b, c and Extended Data Fig. 2a). We validated cluster-specific marker genes *in vivo*. For instance, *in situ* probes against *Vha100-a* identified specifically the copper cell population in the middle midgut, while specific *Gal4* enhancers confirmed *Npcf2* and *dmGlut* expression in various EC populations (Extended Data Fig. 2b-c). We also found that the amino acid transporter *path* marked ISCs, suggesting a potential requirement for nutrient sensing in this cell type (Extended Data Fig. 2b, c), as previously described^14^. Moreover, by mapping bulk regional expression profiles from dissected midguts^30^, we identified the spatial coordinates of all major intestinal cell types (Extended data Fig. 2d, e). In conclusion, our dataset under homeostatic condition provides a comprehensive single-cell atlas of the intestine, offering insights into the regional and functional properties of this organ and serves as a reference map to compare the effects of genetic perturbations.

**Fig. 2.**
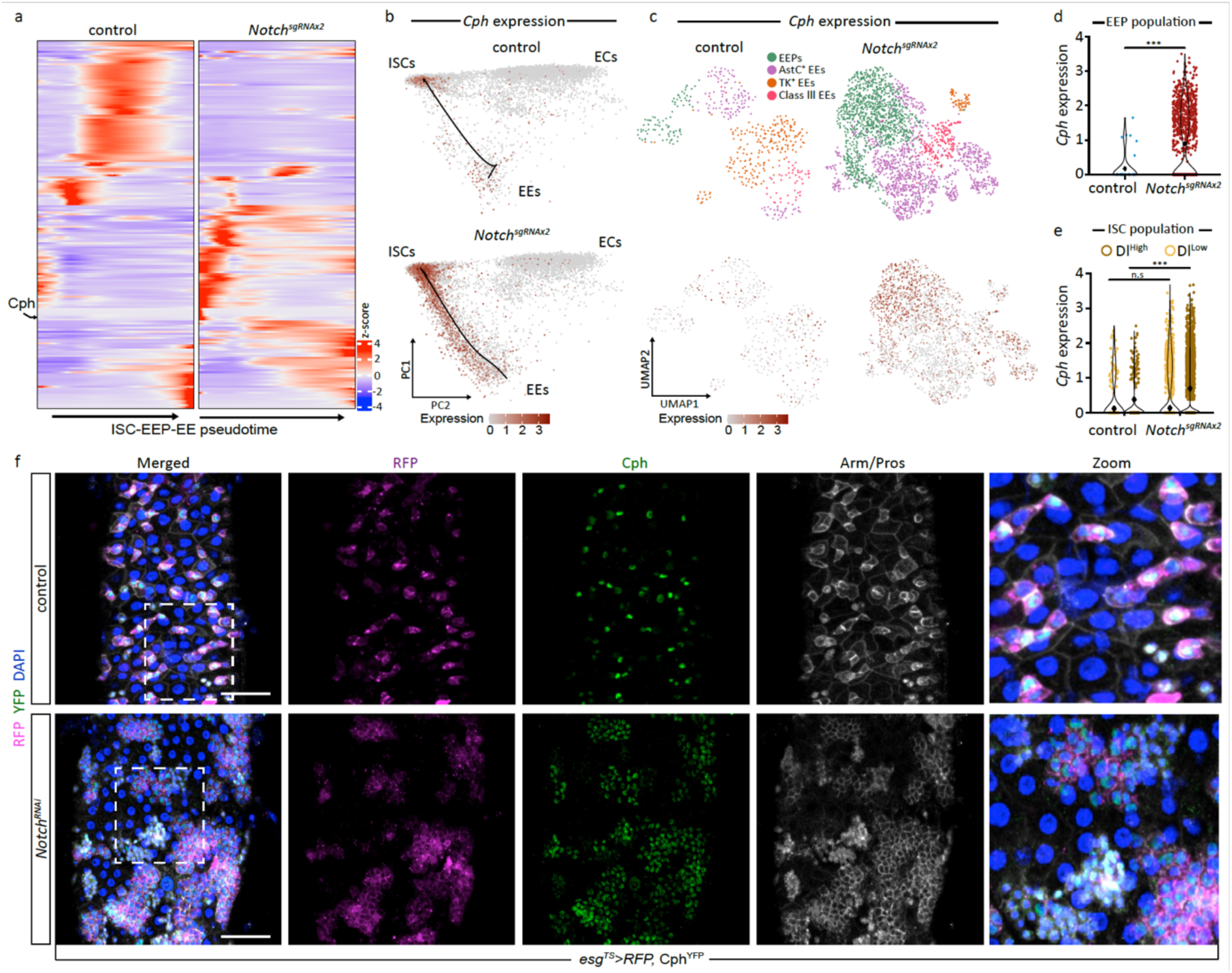
Notch signalling negatively regulates *Cph* expression along the ISC-EE lineage. **(a)** Pseudotime expression analysis of all differentially expressed genes along the ISC-EEP-EE lineage under control and *Notch* mutant conditions. **(b)** PCA projection displaying *Cph* expression in control and *Notch* mutant condition. Notice that *Cph* expression increases specifically along the ISC-EE lineage when *Notch* is mutated. **(c)** *Cph* expression in the EEP and EE population in control and *Notch* mutant condition. *Cph* is present in some EEs during homeostasis and its expression increases mainly in the EEP population. **(d)** Log counts for *Cph* expression specifically within the EEP population under control and *Notch* mutant condition. **(e)** Log counts for *Cph* expression within ISCs that are sub-clustered based on Dl^High^ and Dl^Low^ expression (see M&M). Notice that *Cph* expression increases in Dl^High^ ISCs when Notch signalling is perturbed. **(f)** Endogenous expression of Cph is present within RFP^+^ progenitor cells as well as EEs marked by Pros. Knockdown of *Notch* results in the expansion of *Cph^+^* progenitor cells and to a lesser extent *Cph^+^* EE cells. Scale bar for f is 100 μm.

Next, we quantified the abundances of all major cell types during homeostasis and under progenitor-specific *Notch^sgRNAx^*^2^ mutagenesis. We observed a drastic expansion of Dl^+^ ISCs, pros^+^ EEs and pros^+^/hdc^+^/esg^+^/Dl^+^ EEPs when *Notch* was deactivated (Fig. 1d). We verified these findings by observing an increase in ISC proliferation and EE generation in progenitor-specific *Notch* mutant flies (Fig. 1e, f). We also found that the EB population was significantly decreased and there was a trend towards a smaller EC population, providing further evidence that loss of Notch activity in the progenitor population favours ISC to EE differentiation (Fig. 1d).

After observing major changes in the EE populations, we sought to explore this in more detail, subcategorising EEs based on the expression of different neuropeptides and their spatial coordinates. We found that loss of Notch signalling drastically increased the class I AstC^+^ EEs in the R5 region, while simultaneously decreasing class II Tk^+^ EEs in the same region (Fig. 1d and Extended Data Fig 2e). There was no change in the fraction of total class III EEs that were positive for CCHa2, but variation across different regions were observed (Extended Data Fig 2e. We validated these findings *in vivo*, by performing immunostaining for AstC^+^ and TK^+^ EEs and confirmed that perturbation of Notch signalling mainly affected the balance between class I and class II EE subtypes (Fig.1e, f).

### Notch inactivation causes significant transcriptional and functional changes in progenitor cells

CRISPR mutagenesis in cell populations often gives rise to genetic mosaics that contain cells with functional alleles of the target gene. To assess the efficiency of *Notch^sgRNAx^*^2^ CRISPR mutagenesis in progenitor cells at single cell resolution, we used MELD along with vertex frequency clustering (VFC)^31^. This method aims to find changes in the probability density of cell states between the control and perturbed condition to identify cell populations affected by the perturbation. Using MELD on the *Notch*^sgRNAx2^ perturbed dataset, we observed that 89.4% of progenitor cells were assigned as perturbed and 10.6% as unperturbed (Extended Data Fig. 3a). We confirmed CRISPR editing using Sanger sequencing, demonstrating expected cut site at one of the *sgRNA* targeting location (Extended Data Fig. 3b). Interestingly, we detected no change in the mRNA expression level of mutant *Notch* in the perturbed cells, an observation that suggests CRISPR-induced edits impact Notch protein function rather than mRNA levels (Extended Data Fig. 3c). To assess the state of Notch signalling, we characterised previously validated target genes that report on Notch activity using our scRNA-seq dataset, including *E(spl)mα-BFM*, *E(spl)m3-HLH*, *E(spl)mβ-HLH* and *Klu*. All tested target genes showed a significant reduction in their expression in the *Notch*^sgRNAx2^ perturbed progenitor population (Extended Data Fig. 3d). These results indicate that expression of *Notch*^sgRNAx2^ elicits loss of Notch activity in the majority of progenitor cells.

To understand transcriptional changes in the progenitor population we performed differential gene expression analysis by comparing perturbed to unperturbed cells as inferred by MELD. This analysis revealed 676 upregulated and 784 downregulated genes in the *Notch^sgRNAx^*^2^ perturbed progenitor population. Similar results were obtained when directly comparing cells from the *Notch^sgRNAx^*^2^ condition with those from the control, disregarding the perturbation’s success, which highlights the perturbation efficiency (Extended Data Fig. 4a). We identified only 9 differentially expressed genes (DEGs) in the unperturbed progenitor population from the *Notch^sgRNAx^*^2^ condition when compared to control, consistent with limited transcriptional changes in these cells (Extended Data Fig. 4b). In the perturbed population, we detected an increase in expression of a number of genes previously reported to be upregulated upon loss of *Notch*, including *scute*, *phyl*, and *asense*, and further identified genes related to the cell cycle, reflecting the increased mitosis observed *in vivo* (Extended Data Fig. 4c). Progenitor-specific upregulated genes were significantly enriched in gene sets related to DNA replication, cell cycle, homologous recombination, nucleotide excision repair and mismatch repair, while downregulated genes were associated with longevity regulating pathways, hypoxia and Notch signalling (Extended Data Fig. 4d). Interestingly, we found that E2F and Myc target genes were significantly increased in *Notch* perturbed progenitor cells, indicating that this may be the main driver of proliferation in progenitor cells (Extended Data Fig. 4d). Since CRISPR mutations are heritable after differentiation, we investigated how Notch mutation elicits transcriptional changes in all major intestinal cell types. We identified that the majority of DEGs are specific to one cell type, demonstrating that *Notch* mutations result in cell type specific transcriptional responses (Extended Data Fig. 4e). Moreover, the majority of DEGs were identified in progenitors and EEs, indicating that these cell types are particularly responsive to the loss of Notch signalling (Extended Data Fig. 4e).

### Cph expression along the ISC-EE lineage is negatively regulated by Notch signalling

TFs regulate precise spatiotemporal transcriptional changes in ISCs to govern differentiation. Since we observed major changes in the abundance of EEs upon expression of *Notch^sgRNAx^*^2^, we performed differential gene expression analysis along the ISC-EEP-EE differentiation trajectory, identifying 179 deregulated genes, of which 12 were TFs (Fig. 2a). One particular TF that had a distinctive expression profile was the C2H2 zinc-finger TF Chronophage (*Cph -* CG9650) (Fig. 2a). *Cph* is homologous to mammalian *BCL11a* and *BCL11b*, which are implicated in acute myeloid leukaemia (AML)^32^, haemoglobin switching^33,34^ and intellectual disability disorder^35^. Genetic screens in the past have implicated *Cph* in regulating Notch signalling in the developing *Drosophila* eye^36^. Recently, *Cph* was reported to be involved in the timing of neuronal stem cell differentiation^37^ during development and locomotor behaviour^38^ but its function in adult somatic stem cells are yet to be described. Under control conditions, *Cph* expression was predominately in ISCs and some EEs, whereas upon loss of Notch signalling, *Cph* expression increased specifically in ISCs and EEPs and not along the ISC-EB-EC lineage (Fig. 2b and Extended Data Fig. 5a-c). Indeed, of all the differentially expressed TFs, *Cph* was the only one to display this expression profile (examples of two other TFs are provided in Extended data Fig. 5d, e). Next, we sought to investigate the characteristics of *Cph* expressing cells. We observed that loss of Notch signalling induces *Cph* expression in Dl^+^ Pros^+^ immature EEPs and not IA-2^+^ Pros^+^ mature EEs expressing AstC or Tk (Fig. 2c, d). By grouping ISCs into either Dl^high^ and Dl^low^ populations we observed that the expression of *Cph* was largely in Dl^high^ ISCs and increased specifically in this population upon loss of Notch signalling (Fig. 2e). Interestingly, Dl^high^ ISCs had a greater differentiation potential than Dl^low^ ISCs, indicating that the former cells have a tendency to differentiate into EEPs (Extended data Fig. 5f). Using an endogenously tagged Cph^YFP^ protein trap line, we confirmed Cph^YFP^ expression in ISCs and some EEs *in vivo* (Fig. 2f and Extended Data Fig. 5g). Under conditional inactivation of *Notch* in progenitors, we observed an expansion of Cph^YFP^ cells in ISCs and EEs (Fig. 2f). Thus, *Cph* expression is induced early along the ISC-EE lineage when Notch signalling is perturbed.

Next, we aimed to explore whether the link between Notch signalling and *Cph* is conserved across species. We found that the zinc finger binding domains of *Cph* are well-preserved across diverse species, including in its mammalian counterparts *BCL11a* and *BCL11b* (Extended Data Fig. 6a). We then examined whether *BCL11a* expression correlates with Notch target genes in AML, noting that *BCL11b* is expressed at low levels in this disease and was not included in our analysis^39^. Investigating transcriptomic data from AML patients^39^ revealed that primitive leukemic blasts have high levels of *BCL11a* but low levels of *NOTCH* receptors and NOTCH target genes (Extended Data Fig. 6b) – mirroring our findings in the fly intestine. These findings suggest that the interaction between Notch signalling and *BCL11a*/*Cph* may indeed be conserved in human and more generally in mammals.

### Cph is required during low Notch signalling to maintain epithelial turnover, EE identity and longevity

The fly intestine, like its mammalian counterpart, is continually renewed in order to maintain homeostasis. To characterise the role of *Cph* during epithelial turnover, we first used the ReDDM lineage tracing system^40^, which labels progenitors with a short-lived mCD8-GFP and long-lived H2B-RFP and their differentiated cells in H2B-RFP only (Fig. 3a). Knockdown of *Cph* (*Cph^RNAi^*) decreased the number of GFP^+^/RFP^+^ progenitor cells, some of which were abnormally large in size with distinctive morphology (Fig. 3b and Extended Data Fig. 7a). Using an independent lineage tracing tool termed *esg^F/O^*, which uses temperature-dependent FLPase expression to constitutively activate an Act>STOP>Gal4 driver by removing the STOP cassette located in between FRT sites^41^, we confirmed that loss of *Cph* decreases progenitor and EE turnover (Extended Data Fig. 7b), indicating that *Cph* is required to maintain the rate of normal epithelial turnover. Since *Cph* expression is significantly increased in *Notch* mutant condition we sought to find out whether silencing *Cph* could rescue the excess proliferation and differentiation phenotypes associated with inactivation of Notch signalling. *RNAi*-mediated depletion of *Notch* resulted in a significant increase in GFP^+^/RFP^+^ progenitor cells as well as RFP^+^ only cells with small nuclei resembling EEs (Fig. 3b). Interestingly, simultaneous loss of *Notch* and *Cph* curbed excess progenitor proliferation and EE turnover, suggesting that *Cph* is required when Notch activity is low to generate new ISCs and EEs (Fig. 3b).

**Fig. 3.**
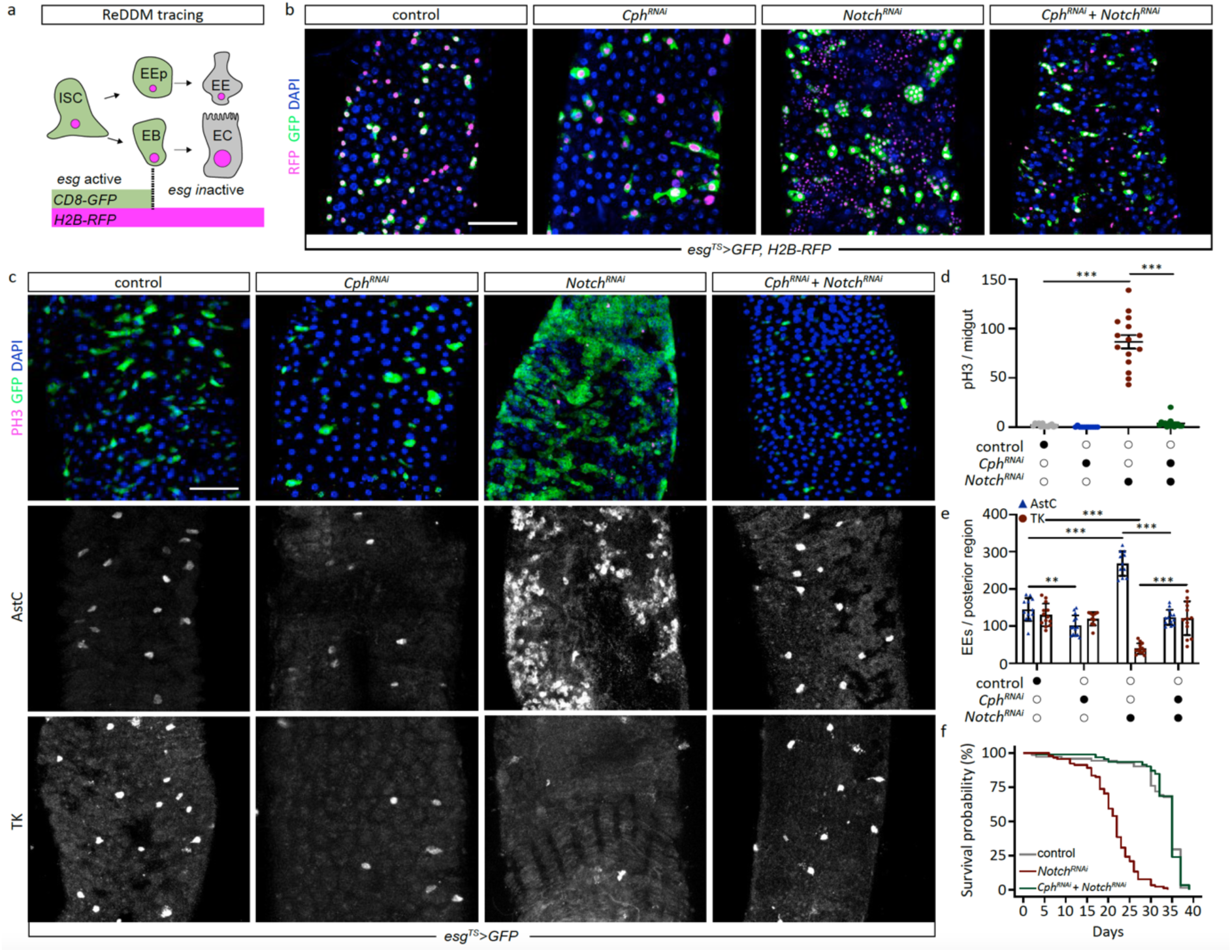
*Cph* maintains and specifies ISCs downstream of Notch. **(a)** Schematic representation of ReDDM lineage tracing tool. ReDDM utilises a short-lived CD8-GFP and a long-lived H2B-RFP. All progenitors were labelled with CD8-GFP and H2B-RFP while the latter was only retained in differentiated cells. **(b)** *Cph* knockdown using the ReDDM system decreases the number of GFP^+^/RFP^+^ cells and moderately increases the number of RFP^+^ only cells. Silencing of *Notch* increases both GFP^+^/RFP^+^ cells and RFP only cells. Co-silencing *Cph* and *Notch* reduces the excess GFP^+^/RFP^+^ cells and RFP only cells as seen with *Notch* depletion only. **(c)** Genetic interaction between *Cph* and *Notch* while monitoring the progenitor population, AstC^+^ EEs and TK^+^ EEs. **(d)** Quantification of mitotic cells. **(e)** Quantification of different population of EEs. **(f)** Quantification of fly lifespan. One-way ANOVA test with Tukey post hoc comparison was used for d and e. \**P* < 0.05, \*\**P* < 0.01, \*\*\**P* < 0.001. A logrank test was used for f. Scale bar for all is 100 μm.

Next, we investigated the function of *Cph* in maintaining different intestinal cell types. We observed that knockdown of *Cph* in progenitor cells significantly decreased the progenitor population as well as AstC^+^ and TK^+^ EEs, suggesting that *Cph* is involved in maintaining the progenitor and EE populations (Fig. 3c-e). As stated above, loss of Notch increased the number of mitotically active ISCs and AstC^+^ EEs while decreasing Tk^+^ EEs. This results in the formation of multi-layered tumour-like structures that consist predominantly of ISCs and EEs which significantly decreases the survival of flies (Fig. 3f). Interestingly, conditional knockdown of both *Cph* and *Notch* decreased the number of progenitors that were mitotically active and resulted in a reversal of EE fate, restoring both AstC^+^ EEs and Tk^+^ EEs back to wild type levels (Fig. 3c-e). Importantly, we were able to also rescue the declining survival of *Notch^RNAi^*expressing flies by co-expressing *Cph^RNAi^* (Fig. 3f). Thus, *Cph* is important for maintaining the progenitor population, EE fate and is physiologically important for longevity under low Notch signalling.

### Scute binds to the *Cph* locus and promotes its expression during low Notch signalling

Previous reports demonstrate that *sc* is transiently expressed in a subpopulation of ISCs in response to low Notch signalling^22^. Sc in turn binds to the *pros* locus, a master regulator of EE fate, to increase its expression and commit ISCs to an EE fate. We confirmed the increase of *sc* expression in the *Notch*^sgRNAx2^ condition (Extended Data Fig. 4c), and further show that this occurs specifically in the Dl^High^ ISC population (Extended Data Fig. 8a). Using our scRNA-seq dataset, we observed an overlap between *sc* and *Cph* in the ISC and EEP populations (Fig. 4a). We further grouped ISCs into *sc^+^*and *sc^-^* populations and found *sc* to be partially co-expressed in *Cph^+^* ISCs (Fig. 4a and Extended Data Fig. 8b). Using a previously validated enhancer trap line that partially recapitulates *sc* expression, we found sparse co-expression between *sc* and *Cph in vivo* (Fig. 4b). Interestingly, *sc^+^* ISCs also displayed low Notch signalling activity and increased expression of cell cycle regulators, including *CycE* and *CycA* when compared to the *sc^-^*populations (Extended Data Fig. 8c). Therefore, *sc* and *Cph* are expressed in a subset of ISCs with low Notch signalling activity.

**Fig. 4.**
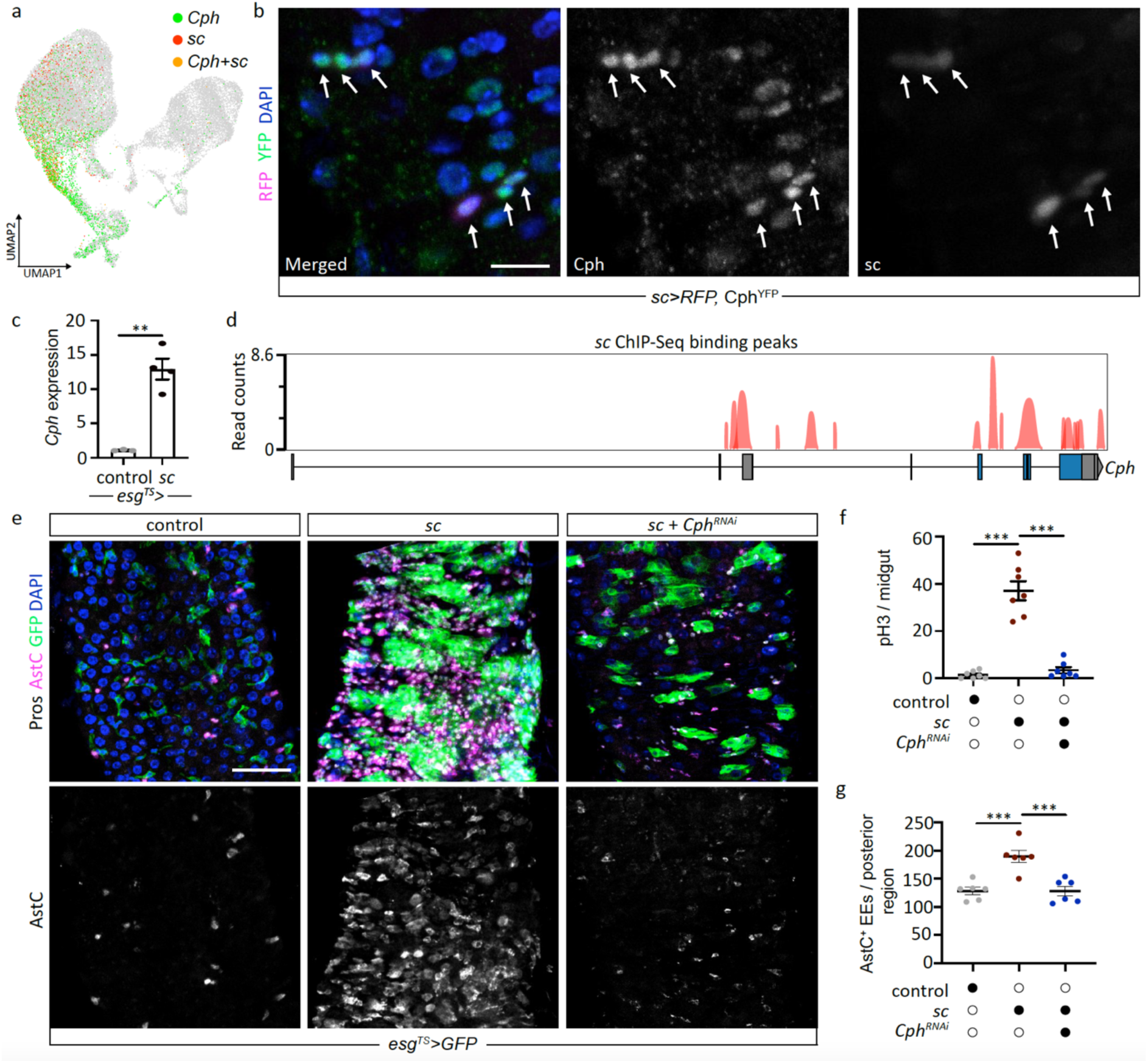
Regulation of *Cph* expression by *scute*. **(a)** UMAP of *Cph*^+^ (green) and *sc*^+^ (red) cells, with co-expression indicated in orange. **(b)** *scute* is co-expressed in a sub-population of *Cph*^+^ cells *in vivo*. **(c)** Bulk RNA-sequencing of scute overexpression in progenitor cells, demonstrating an upregulation in Cph expression. **(d)** Chip-seq binding peaks of scute within the *Cph* locus. Multiple peaks are observed corresponding to the transcriptional start site (TSS) of Cph isoforms. **(e)** Overexpression of *sc* in progenitor cells increases the number of progenitors as well as AstC^+^ EEs. Silencing *Cph* in progenitor cells overexpression *sc* suppresses the phenotypes associated with *sc* overexpression. **(f)** Quantification of mitotic cells. **(g)** Quantification of AstC^+^ EE cells. One-way ANOVA test with Tukey post hoc comparison was used for F and G. \**P* < 0.05, \*\**P* < 0.01, \*\*\**P* < 0.001. Scale bar for b is 10 μm and for e is 100 μm.

Since we observed an overlap in the expression of *sc* and *Cph,* we investigated whether Sc is able to directly regulate *Cph* expression. By re-analysing previously published bulk RNA-sequencing data from intestinal progenitor cells overexpressing *sc*^22^, we found *Cph* expression to be significantly upregulated in this condition (Fig. 4c). To gain mechanistic insight into how *sc* regulates *Cph* expression, we surveyed publicly available ChIP-seq data obtained for Sc in the progenitor population^42^. We confirmed previous reports that Sc binds to the *pros* locus and further demonstrated the specificity of the ChIP-seq data by showing that Sc does not bind to *rdhB*, which is enriched in the adult eye and not expressed in the intestine (Extended Data Fig. 8d). Interestingly, we detected Sc binding peaks at *Cph* transcriptional start sites (TSS), specifically in the TSS region of the transcripts Cph-Rl and Cph-RK/Cph-PL and on the exons of Cph-RK/Cph-PL (Fig. 4d). In support of this, we also found the E-box binding motif GCAGGTGT within the *Cph* locus, which has previously been shown to be bound by *sc*^43^ (data not shown). These data suggest that Sc can directly regulate *Cph* expression.

In light of the findings above, we hypothesised that *Cph* may function downstream of *sc* to regulate progenitor proliferation and the generation of EEs. We observed that *sc* overexpression in progenitor cells resulted in a significant expansion of progenitor cells that were actively dividing and in the number of EE cells positive for AstC (Fig. 4e-g). Knockdown of *Cph* in progenitor cells overexpressing *sc* rescued both of these phenotypes, indicating that *Cph* lies downstream of *sc* (Fig. 4e-g). In conclusion, our findings suggest that during low Notch signalling *sc* directly binds to the *Cph* locus to induce its expression along the ISC-EE trajectory.

### Cph is required to remodel the transcriptome of progenitor cells upon loss of Notch signalling

Thus far, we have shown that when Notch signalling is inactivated *Cph* expression is elevated along the ISC-EE lineage and is required for maintaining the progenitor and EE populations. To investigate the transcriptional programs that *Cph* regulates during this process, we performed scRNA-seq of the intestine while silencing either *Notch^RNAi^* or *Cph^RNAi^+Notch^RNAi^*in the progenitor population. After quality control, we recovered 16,750 cells across control and perturbed conditions and identified all major cell type clusters (Fig. 5a). We confirmed that both *Notch* and *Cph* expression were downregulated in ISCs, demonstrating the effectiveness of *RNAi* silencing (Extended Data Fig. 9a, b). We also show that the differentially expressed genes between *Notch^RNAi^* and *Notch*^sgRNAx2^ conditions were positively correlated, indicating that both perturbations elicit similar transcriptional changes within the progenitor population (Extended Data Fig. 9c). Indeed, a notable increase in *Cph* expression was also observed in progenitor cells expressing *Notch^RNAi^* (Extended Data Fig. 9b), underscoring the strong induction of *Cph* expression when Notch signalling is disrupted, whether through *RNAi* silencing or CRISPR mutagenesis.

To investigate changes in epithelial cell composition, we quantified cell type abundances across all conditions. We observed a notable increase in the number of ISCs, EEPs, EEs and a decrease in EBs in the *Notch^RNAi^* condition (Fig. 5b), consistent with observations made using CRISPR-Cas9 (Fig. 1c). Interestingly, when progenitor cells expressed *Cph^RNAi^+Notch^RNAi^*, the number of ISCs, EEPs and EEs decreased in comparison to *Notch* silencing (Fig. 5b). Furthermore, while the number of ECs were lower when *Notch* was depleted, co-silencing *Cph^RNAi^+Notch^RNAi^* resulted in a significant increase of ECs, although this was not accompanied by an increase in the EB population (Fig. 5b). These data suggest that *Cph* is required when Notch signalling is low to maintain different intestinal cell types.

To gain mechanistic insight into the differences in cell type composition, we identified DEGs in specific progenitor populations, including ISCs only, ISCs and EBs, and ISCs and EEPs. We identified multiple Notch target genes, such as *E(spl)mβ-HLH* and *E(spl)mα-BFM*, which were significantly downregulated when *Notch* was silenced, indicating a decrease in Notch signalling activity (Fig. 5c). We also observed changes in the expression of key TFs involved in ISC differentiation, including *sc* and *klu* (Fig. 5c). Moreover, genes related to the cell cycle and ISC maintenance were significantly upregulated in the *Notch^RNAi^* condition, including *CycA*, *mira* and *Dl* (Fig. 5c). Interestingly, many of these genes were downregulated in progenitors (ISCs and EEPs) expressing *Cph^RNAi^+Notch^RNAi^*, suggesting that *Cph* is involved in inducing their expression (Fig. 5c). A quantitative display of the intersection between the number of differentially expressed genes highlighted that the third and fourth largest groups belonged to the shared deregulated genes in *Notch^RNAi^* and *Cph^RNAi^+ Notch^RNAi^* (Fig. 5d). To more directly understand the requirement of *Cph* to remodel the transcriptome of *Notch* depleted progenitor cells, we used Pearson’s correlation coefficient to compare the relationship between shared deregulated genes in *Notch^RNAi^* and *Cph^RNAi^+ Notch^RNAi^* condition. This revealed that the transcriptional signatures of *Notch* depleted ISCs and ISC+EEPs are blunted when *Cph* is silenced (Fig. 5e and Extended data Fig. 9d, e). Moreover, hallmark gene sets analysis^44^ revealed an enrichment of terms related to Myc targets and DNA repair, and a downregulation of terms associated with inflammatory response, NF-κB signalling and JAK-STAT signalling when Notch was silenced. Interestingly, these terms were reversed following co-expression of *Cph^RNAi^+Notch^RNAi^* (Fig. 5f). Importantly, this transcriptional rewiring was not due to increasing *Notch* expression or Notch signalling, as the expression of *Notch* and Notch pathway target genes, such as *E(spl)mβ-HLH* and *E(spl)mα-BFM*, were significantly downregulated in both the *Notch^RNAi^* and *Cph^RNAi^+Notch^RNAi^* conditions (Fig. 5c). Conversely, we observed a positive correlation between the commonly deregulated genes in the ISC+EB population, indicating that *Cph* has limited roles in altering the transcriptome of these progenitors when *Notch* is depleted (Extended data Fig. 9e). In conclusion, our findings suggest that *Cph* is required to reprogram the transcriptional state of ISCs and EEPs under conditions of reduced Notch signalling.

### Chromatin binding profile of Cph in progenitors reveals novel target genes involved in ISC proliferation and EE generation

Next, we sought to identify Cph target genes *in vivo* and performed NanoDam to generate DNA binding profiles for Cph (Extended Data Fig. 10a). NanoDam utilises a GFP-recognising nanobody which links Dam methylase to an endogenously tagged DNA-binding protein, resulting in m6A methylation of surrounding GATC sites^45^. This method is advantageous over other chromatin profiling approaches such as targeted DamID (TaDa)^46^ as it does not require overexpression of DNA-binding protein. We restricted expression of NanoDam specifically to the progenitor population using the *esg^TS^* driver and induced its binding to endogenously tagged Cph-YFP for approximately 17h (Extended Data Fig. 10b). After quality control and normalisation to the NanoDam-only control condition (Extended Data Fig. 10c), significant Cph binding sites were found for 807 genes across the entire genome (Extended Data Fig. 10d). We observed pronounced peak intensity profiles for cell cycle related genes, including *CycE,* and *E2F1* (Fig. 6a). Moreover, we also identified a significant intronic peak in *pros* (Extended Data Fig. 10e), suggesting that *Cph* may regulate its expression. Conversely, we did not identify significant peaks in *rtp*, which is not expressed in the intestine during homeostasis, suggesting that our dataset is robust (Extended Data Fig. 10e).

**Fig. 5.**
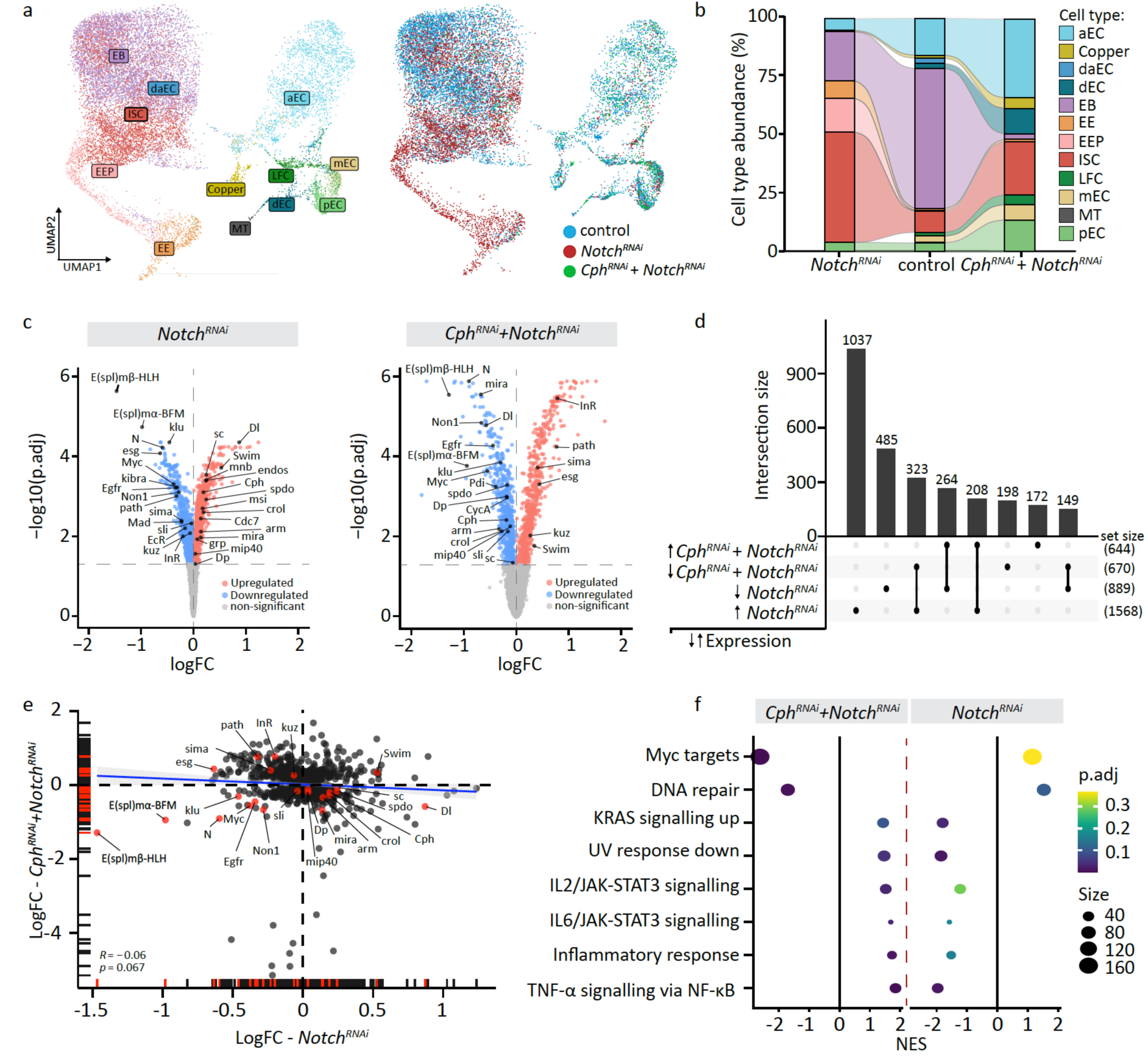
*Cph* is required to remodel the transcriptome of Notch depleted intestinal stem cells. **(a)** UMAP plot coloured by intestinal cell types and split by perturbation conditions. **(b)** Alluvial plot representing changes in cell type abundance in the respective perturbation condition. **(c)** Volcano plot highlighting differentially expressed genes in ISCs and EEPs. **(d)** Upset plot illustrating the number of differentially expressed and their intersect between groups. Arrows indicate whether gene expression is up or down in perturbed condition and total number of genes are provided in set size. **(e)** Pearson’s correlation coefficient of all shared deregulated genes in *Notch^RNAi^*and *Cph^RNAi^+Notch^RNAi^*. **(f)** Hallmark gene sets enrichment analysis results (only the enriched pathways with opposite effects between the two perturbations are shown).

**Fig. 6.**
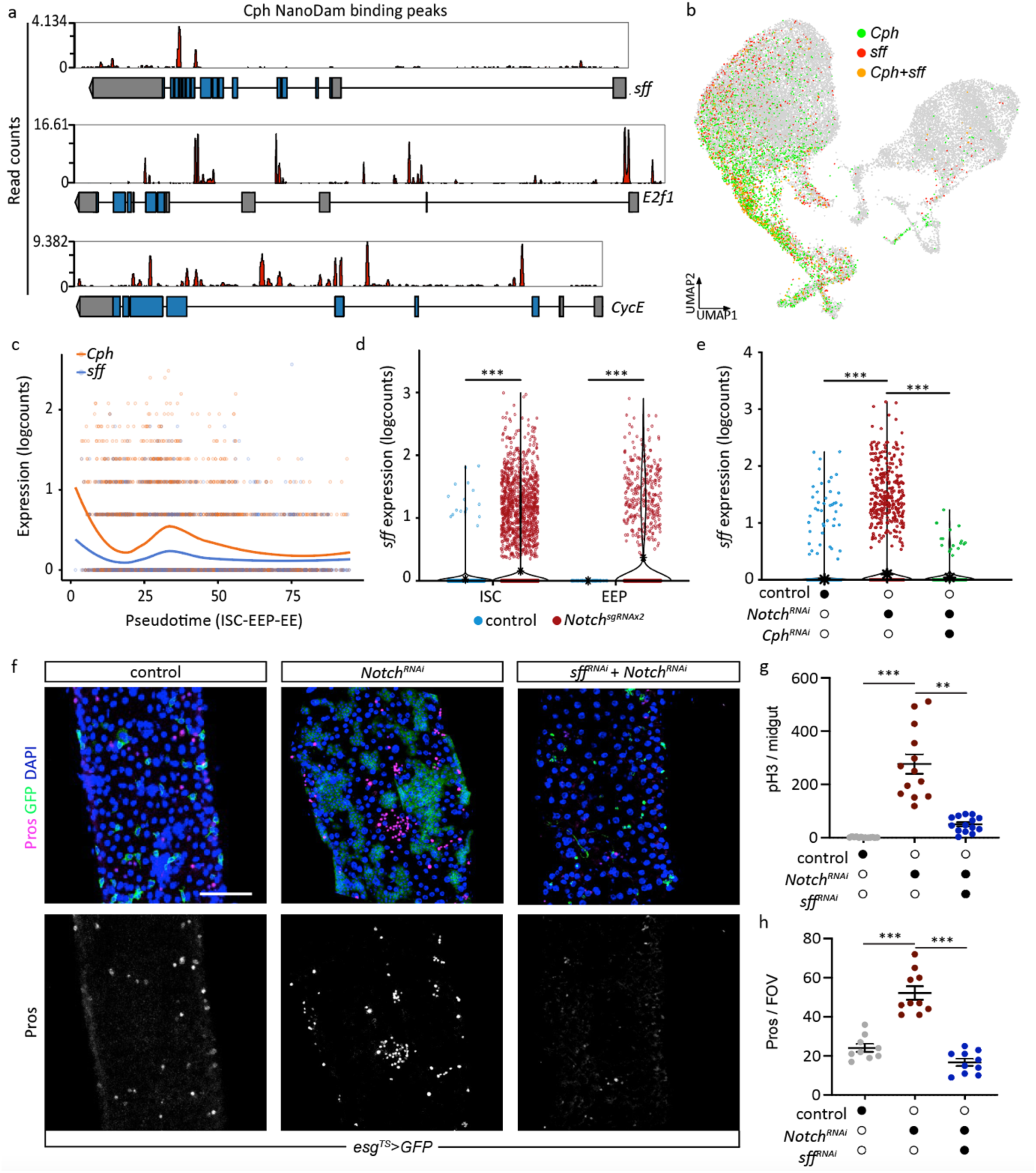
NanoDam identifies *Cph* target genes that important for proliferation and differentiation of ISCs *in vivo*. **(a)** NanoDam binding profile for Cph in various target genes are shown as summed coverage difference relative to NanoDam only. **(b)** UMAP of *Cph*^+^ (green) and *sff*^+^ (red) cells, with co-expression indicated in orange. **(c)** Expression of *Cph* and *sff* along the ISC-EEP-EE lineage. Notice a burst in expression of both *Cph* and *sff* early along this trajectory. **(d)** Expression of *sff* in ISCs and EEPs during homeostasis and Notch mutant condition. **(e)** Expression of *sff* in all progenitors during homeostasis, *Notch^RNAi^*and *Cph^RNAi^+Notch^RNAi^* conditions. **(f)** Knockdown of *sff* in progenitors depleted of Notch significantly reduces GFP+ progenitors and pros+ EEs when compared to *Notch^RNAi^* expressing progenitors. **(g)** Quantification of mitotic cells in the entire midgut. **(h)** Quantification of EEs in the field of view. One-way ANOVA test with Tukey post hoc comparison was used for (g) and (h). \**P* < 0.05, \*\**P* < 0.01, \*\*\**P* < 0.001. Scale bar for f is 100 μm.

One of the top genes identified in our Cph NanoDam dataset was sugar-free frosting (*sff*) (Fig. 6a), which is poorly characterised with respect to its function in the intestine. *sff* was particularly interesting because it was barely detectable during homeostasis but its expression increased in *Notch^sgRNAx^*^2^ mutant flies, coinciding with *Cph* along the ISC-EE trajectory (Fig. 6b, c, d). The induction in *sff* expression was prominent in ISCs and EEPs after Notch signalling was disrupted using either *Notch^sgRNAx^*^2^ or *Notch^RNAi^* (Fig. 6d, e). Interestingly, the expression of *sff* was returned to wild type levels after co-silencing *Cph* and *Notch*, indicating that *Cph* directly regulates the expression of *sff* (Fig. 6e).

*sff* has previously been characterised as an SAD like kinase that is important regulator of vesicle tethering and glycosylation in embryos^47^. Since *sff* expression is induced during low Notch condition, we asked whether it is functionally required for the proliferation and differentiation of *Notch* depleted tumours. While flies expressing progenitor-specific *Notch^RNAi^* developed intestinal neuroendocrine tumour-like structures characterised by excessive progenitors and EE cells (Fig. 6f), silencing *sff* significantly reduced tumour formation (Fig. 6f). Consistently, we also observed a significant decrease in the fraction of mitotically active cells in flies expressing *sff^RNAi^* + *Notch^RNAi^*compared to *Notch^RNAi^* only (Fig. 6g). Moreover, co-silencing of *sff* and *Notch* highlighted a key requirement for *sff* in the formation of EEs (Fig. 6f, h). These findings reveal a previously unrecognised role for vesicle tethering and glycosylation in driving hyperproliferation and differentiation of ISCs. In conclusion, our *in vivo* chromatin binding profiles of Cph highlighted key genes involved in cell cycle regulation and EE differentiation and further revealed a novel role for *sff*, which is required for ISC proliferation and EE generation.

## DISCUSSION

We present a comprehensive multi-omics analysis of the fly intestine combined with single and co-perturbation studies. We identify *Cph* as a novel regulator of ISC maintenance and differentiation downstream of Notch. *Cph* is present in ISCs and EEs during homeostasis and its expression is negatively regulated along the ISC-EE trajectory by Notch signalling. The expression of *Cph* is under the control of *scute*, which binds to multiple sites within the *Cph* locus to promote its expression after Notch signalling is inactivated. *Cph* coordinates proliferation and differentiation of ISCs and EEPs by directly regulating cell cycle genes and *pros*. A brief burst of *Cph* expression specifically during the early stages of ISC-EE differentiation ensures that cell fate commitment is faithfully executed. Our scRNA-seq of the intestine provides an atlas of both homeostatic and *Notch* mutant conditions, including their spatial features. This dataset may also serve as a crucial reference map for future research into the role of Notch signalling in the intestine and can be explored by the community using our Shiny App.

By profiling chromatin binding sites of Cph *in vivo* using NanoDam, we identified hundreds of target genes that are bound by Cph in progenitor cells. Indeed, we identified *sff*, a homolog of SAD kinase, as a crucial *Cph* target gene that regulates ISC proliferation and EE generation. Very little is known about the function of *sff* outside the context of the embryo where it has been described to be involved in vesicle tethering and glycosylation at the Golgi^47^ – processes that are important for membrane trafficking. Given that membrane trafficking and endocytic events are crucial for regulating Notch signalling^48^, our findings raise the possibility that *sff* may also be involved in regulating membrane trafficking of Notch pathway components.

While our findings demonstrate that *sc* can directly regulate *Cph* expression, it is crucial to recognize that *sc* is only transiently expressed in ISCs during homeostasis^22^, whereas *Cph* expression remains stable. This implies that additional transcription factors may be involved in regulating *Cph* expression. Alternatively, *Cph* might regulate its own expression through autoregulation. Similar mechanisms have been proposed for key TFs in the intestine, such as *klu*^24^ and *sc*^22^, which either suppress or enhance their own expression, respectively. Beyond transcriptional regulation, post-translational modifications may also play a role in controlling Cph levels in ISCs. Previous studies have identified Phyl as an adaptor protein that promotes the degradation of Ttk via the E3 ubiquitin ligase pathway^27^. Notably, we observed a significant upregulation of *phly* following Notch signalling inactivation, however, further research is needed to determine whether Phly protein also regulates Cph levels in ISCs.

Cph is homologous to mammalian BCL11a and BCL11b. BCL11a is strongly expressed across various hematopoietic lineages and is involved in the transition from γ-globin to β-globin expression during the shift from fetal to adult erythropoiesis^33,34^. Moreover, BCL11a is well documented to promote acute myeloid leukemia (AML)^32^, while Notch signalling is implicated in the regulation of hematopoietic stem cells^49^. Our analysis of bulk transcriptomic datasets from primitive leukemic blasts highlighted that BCL11a and Notch signalling target genes are anticorrelated – when BCL11a expression is high, Notch target genes are low, mirroring our observation in the fly intestine. This suggests that Notch signalling and BCL11a/Cph may have conserved roles in regulating adult stem/progenitor cell types in other tissues and may represent novel therapeutics targets to combat human cancers.

## MATERIALS AND METHODS

### Fly husbandry

*Drosophila* stocks were raised on a 12:12 hour light:dark cycle and maintained on standard fly food consisting per litre of 44 g syrup, 80 g malt, 80 g corn flour, 10 g soy flour 18 g yeast, 2.4 g methly-4-hyroxybenzoate, 6.6 mL propionic acid, 0.66 mL phosphoric acid and 8 g agar. For CRISPR mutagenesis experiments, newly eclosed flies were shifted to 29°C for 10 days and subsequently to 18°C for 30 days before switching back to 29°C for one day prior to imaging as previously reported^50^. For all other experiments requiring temperature shift (*Gal80^TS^*) for transgene induction, parental lines were kept at 18°C and the progeny were shifted to 29°C after eclosion for 20 days. For all experiments, mated female flies were used and transferred to fresh food every two days.

### Fly stocks

The following fly lines were used in this study: *esg^TS^, UAS-GFP, UAS-Cas9*^p.2^ (gift from F. Port)*, esg^TS^, UAS-GFP* and *esg^TS^, UAS-RFP* (gift from B. Edgar)*, esg^TS^*(gift from B. Edgar), *UAS-H2B-RFP* (gift from Tobias Reiff^40^)*, esg^F/^*^O^ (gift from P. Patel)*, Pros^TS^* (Boutros lab), *dmGlut-Gal4* (BL 63397), *Npc2f-Gal4* (BL 81176), *Path-Gal4* (BL 71411), Cph-YFP (gift from Andrea Brand, DGGR 115236), *UAS-NanoDam^RFP^* (Gift from Andrea Brand^45^) *UAS-Cph^RNAi^* (VDRC 104402), *UAS-Cph^RNAi^ ^#2^* (BL 26713), *UAS-Notch^RNAi^* (VDRC 27228), *UAS-Notch^sgRNAx2^* (VDRC 341922), *UAS-sc* (BL 51672), *UAS-sff^RNAi^* (VDRC 100717).

### Escargot-Flip-Out experiments

Flip-out clones were generated as previously described. Briefly, expression of *flippase* by esg^TS^ at 29°C activates a constitutive Act>STOP>Gal4 driver by excising the STOP cassette flanked by FRT sites. This system was induced for 15-20 days and results in expression of *GFP* and *RNAi* in both progenitor cells and their descendant progeny. Flies were evenly housed in control and treatment groups and their food was changed every 2 days throughout the experiment.

### Dissection and immunohistochemistry

Flies were starved for 3h prior to dissection to reduce luminal content in the intestine. Adult female intestines were dissected in PBS (Phosphate buffered saline, P3812-10PAK) and transferred to Polylysine slides and fixed in 4% Paraformaldehyde (16 % Paraformaldehyde (Thermo Scientific) diluted in PBS) for 20-60 min depending on the antibody. Samples were washed with PBST (PBS with 0.1% Triton X-100) for 30 min and then blocked with PBSTB (PBST with 1% Bovine serum albumin) for 30 min at room temperature. Primary antibody was diluted in PBSTB and incubated with samples overnight at 4°C. Samples were then washed five times in PBST and incubated for 1.5 h-2.5h at room temperature with secondary antibody (antibodies coupled to Alexa fluorophores, Invitrogen) in PBSTB. Samples were washed five times in PBST and mounted in mounting medium (VECTASHIELD from Vector Laboratories with or without DAPI; Vector Labs, H-1200 or H-1000 respectively). Immunostainings for both experimental and control conditions were carried out on the same slide to enable direct comparisons. The following antibodies were used: mouse anti-Armadillo (1:50; DSHB, N27A1), rabbit anti-GFP (1:1000; Invitrogen, A-11122), mouse anti-Prospero (1:20; DSHB, MR1A), anti-Tachykinin (1:500, Gift from Jan A Veenstra^51^), anti-Allatostatin C (1:500, Gift from Jan A Veenstra^51^), rabbit anti-Phospho-Histone H3 (1:500; Cell Signaling, 9701S). Conjugated fluorescent secondary antibodies (conjugated to AlexaFluor488, AlexaFluor549 and AlexaFluor549) were obtained from Invitrogen (Life Technologies) and used at 1:1000.

### Fluorescent *In situ* hybridisation

The following primer pair were used to generate *Vha100-4* probes; F: GAGAGCAACAGCATCTTCCG, R: CAGCACTTGGATCATCTCGC. Probes were labelled using a DIG RNA labelling mix (Roche) following the manufactures guidelines. Intestinal samples were prepared and fixed as previously reported^52^.

### Image acquisition and processing

Confocal fluorescent images were acquired using either an upright Nikon A1 confocal microscope with a 25x Apo dipping objective (NA 1.1) or a spinning disk microscope (CREST V3) on a Nikon Ti2 inverted microscope equipped with a 60x NA PlanApo 1.4 oil immersion objective using Nis-Elements 5.3 software. Images shown represent maximal intensity projection of stacks covering the first epithelial layer. The same acquisition settings (laser power and gain/ camera settings) were applied to both experimental and control groups. All statistical analyses were performed on raw 16bit images using Fiji 2.0 (see detailed description below).

### Lifespan assay

10-15 adult female flies and five adult male flies were kept in each vial as described in Fly husbandry at 29°C. All lifespan experiments started with at least 50 adult female flies for each condition and the number of dead female flies was recorded every day until all female flies were dead.

### Single cell RNA-sequencing and high-throughput sequencing

Flies were starved for 3h and placed in vials with filter paper containing 5% sucrose for 16h. 20 midguts were dissected from respective genotypes in ice-cold PBS, taking care to remove the hindgut, Malpighian tubules and proventriculus. Samples were digested in 1 mg /ml Elastase (Sigma, #E0258) solution at 25°C for 45 min at 1000RPM and vortexed every 15min. Samples were then pelleted and resuspended in PBS and dissociated cells were then passed through a 40uM than 20uM cell strainer and subsequently counted. Approximately 20,000 live cells were used for scRNA-seq with 10X Genomics using either the 3’ kit for CRISPR experiments or 5’ kit for RNAi experiments following the manufacturers protocol for library generation. Prior to sequencing, library fragment size was determined using an Agilent Bioanalyzer high-sensitivity chip and quantified using Qubit. Libraries were multiplexed and sequenced using a Nextseq 550 at the Deep Sequencing Facility, BioQuant, Heidelberg University.

### Single-cell RNA-sequencing data analysis

A CellRanger (version 7.0.1) index was built using the *Drosophila melanogaster* genome sequence along with the corresponding GTF file (Ensembl release 102). To generate single-cell count matrices, reads were aligned to the reference using *cellranger count* with the *include-introns* option. Single-cell RNA-seq data quality control and analysis were conducted using the R package Seurat (version 4.3.0). Raw gene expression count matrices were pre-processed and filtered using scran^53^ (version 1.24.0), following the guidelines of Amezquita et al^54^. In brief, droplets with RNA content below each library’s inflection point were removed, along with cells meeting any of the following criteria: (a) fewer than 250 detected genes, (b) ranking in the top 1 percentile of cells by UMI count, or (c) a high percentage of mitochondrial reads. One replicate of the *Notch^RNAi^* and *Cph^RNAi^*+*Notch^RNAi^*condition showed an unusually high content of mitochondrial reads and were removed from subsequent analyses. Seurat (version 4.1.1)^55^ was used for all downstream analyses unless otherwise stated. These analyses included, log normalization of counts, data scaling, cell cycle inference, and identifying the 3000 most variable genes per replicate. Afterwards the replicates were integrated and batch effects were corrected using the IntegrateData approach from Seurat. Dimensionality reduction was performed using RunPCA and UMAPs were constructed using the first 20 principal components. Clustering of the data was performed using the Louvain algorithm^56^. Cell type labels were manually assigned to clusters based on the expression of characteristic marker genes^14,16,30^. Subpopulations of EEs were identified by reclustering EEs and analysing marker gene expression^16^. For differential cell type abundance, we quantified the number of cells per cell type and replicate and modelling the resulting data as a negative binomial distribution using DESeq2 (version 1.36.0)^57^. The size factors were set to be equal to the total number of cells per replicate.

### Regional mapping of intestinal cells

We downloaded the summary table from region-specific bulk RNA sequencing dataset^30^ to predict the regional origin of our cells. Using SingleR (v1.10.0)^58^ with default settings, predictions were performed individually for ISCs, EBs, ECs, and EEs.

### Trajectory differential expression analysis

To reduce the complexity of the dataset for the trajectory inference, we aggregated large flat cells (LFCs) and copper cells into middle enterocytes (mECs) and differentiating anterior enterocytes (daECs) into differentiating enterocytes (dECs). Using 30 principal components as the input, individual trajectories for the control and *Notch^sgRNAx2^*condition were then inferred using *slingshot* (version 2.4.0)^59^ and condiments (version 1.6.0)^60^, with ISCs as the starting cluster. The associationTest function from tradeSeq (version 1.13.04)^61^ was used to determine gene expression changes along a differentiation trajectory, while the conditionTest function was used to identify condition-specific gene expression changes. Gene expression trends along trajectories were generated by local polynomial regression fitting as implemented by the *loess* function in R.

### MELD analysis

Knockout efficiency was assessed independently for each replicate using MELD (version 1.0.0)^31^, which calculates the likelihood that a particular cell is from the control or *Notch^sgRNAx2^* condition. To identify the optimal set of parameters for the MELD algorithm, we performed a parameter search using the Benchmarker class as implemented in the MELD package. Ultimately, the algorithm was run using 20 principal components as the input. Subsequently, for each cell type, the determined perturbation likelihoods were used as inputs for vertex frequency clustering (VFC)^31^ to identify a population of perturbed and unperturbed cells respectively.

### Differential expression and gene set enrichment analysis

Where possible, pseudobulk expression profiles were created by aggregating counts of the same cell type across replicates and perturbation status, as inferred by MELD. Subsequently, expression was compared using *muscat* (version 1.12.1)^62^ and *edgeR* (version 3.38.0)^63^. For experimental conditions with a single replicate, we applied a generalized linear mixed model with a random effect in accordance with Zimmerman et al.’s^64^ recommendations using MAST^65^ or Seurat’s FindMarkers along with the *test.use=”MAST”*parameter Gene set enrichment analysis was performed using *fgesa* (version 1.22.0)^66^. Gene sets were taken from the FlyEnrichR website (https://maayanlab.cloud/FlyEnrichr/#stats) and the msigdbr package (version 7.5.1).

### Single cell differentiation potential

To assess single-cell differentiation potency, we employed *CompCCAT* from the R package SCENT (version 1.0.3). The necessary protein-protein interaction network was downloaded from the Molecular Interaction Search Tool (https://fgrtools.hms.harvard.edu/MIST/downloads.jsp).

### Categorizing expression levels

To classify cells based on gene expression levels, we computed the 25^th^ and 75^th^ percentiles for each gene and condition. Cells at or below the 25^th^ percentile were categorized as having “low” expression, while those at or above the 75^th^ percentile were classified as “high.” Additionally, if the 75^th^ percentile of expression was less than 1, any cell with an expression level equal or greater than 1 was classified as “high.”

### Bulk RNA-seq Data Analysis

Processed bulk RNA-seq data was downloaded from GEO (GSE102569, Source Paper DOI: 10.1038/s41556-017-0020-0). We removed genes with less than 10 counts in total. Next, we used PyDESeq2 (v0.4.9, DOI:10.1093/bioinformatics/btad547) for differential gene expression testing between control and *Sc* overexpressed samples using a Wald-test. Gene *Cph* (CG9650) showed a p-value of 0.000097 for upregulation under the *Sc* overexpressed condition.

### CHIP-seq and DamID Data Analysis

Raw files were acquired via sratools (v3.0.10) from the respective GEO repositories (CHIP-seq: GSE84283 and DOI: 10.1038/s41598-017-01138-z; DamID: GSE211629 and DOI: 10.1038/s41467-022-34270-0) and extracted as fastq files. Reads were trimmed using trim_galore (v0.6.10, DOI: 10.5281/zenodo.7598955) with settings for illumina reads, removing 10bp from both the 5’ and 3’ ends of all reads, and requiring a minimal read length of 36bp. Trimmed reads were then aligned to the *Drosophila Melanogaster* genome using a precompiled bowtie index (BDGP6) and bowtie2 (v2.5.4, DOI:10.1038/nmeth.1923) while allowing for maximum single mismatches. Resulting alignment files were filtered for uniquely mapped reads, converted to bam, sorted, and indexed using samtools (v1.20, DOI:10.1093/bioinformatics/btp352). We used MACS2 (v2.2.9.1, DOI:10.1101/496521) for peak calling while filtering out peaks with FDR>0.05. Peaks were then annotated with CHIPseeker (v1.38.0, DOI:10.1093/bioinformatics/btv145) and a genome annotation file from Ensembl (index BDGP6.46.110). Tracks were plotted using pyGenomeTracks do:10.1038/s41467-017-02525-w, di: 10.1093/bioinformatics/btaa692. The entire workflow was implemented using snakemake (v8.14.0, DI: 10.12688/f1000research.29032.1).

### Cph NanoDam

To perform Cph NanoDam in the intestine, we crossed a Cph^YFP^; *UAS*-*NanoDam* line with *esg^TS^*. As control, we crossed *UAS*-*NanoDam* line with *esg^TS^*. Flies were reared at 18°C. Adult flies aged between 3-5 days were shifted to 29°C for 17hrs. 30 Midguts where dissected in ice-cold PBS, taking care to remove Malpighian tubules, hindgut and proventriculus and samples were frozen in -80°C. NanoDam samples were processed as previously described^45^. DNA was extracted from dissected midgut and methylated fragments were isolated with DpnI and DpnII digestion. Methylated fragments were then amplified with PCR and sonicated in order to generate libraries suitable for sequencing. Sequencing was performed using single end 86 bp reads with a using a Nextseq 550 at the Deep Sequencing Facility, BioQuant, Heidelberg University.

### NanoDam Data Analysis

The reads were processed as described for the CHIP-seq data. Additionally, a GATC signal was obtained by binning the genome into consecutive GATC sites using a custom python script and a GFF of sites obtained from damidseq_pipeline doi:10.1093/bioinformatics/btv386, then calculating fold-changes of the NanoDam Cph signal with respect to the control for each replicate respectively.

### Statistics

GraphPad Prism 10 software was used for statistical analyses. The statistical tests used for each experiment are indicated in the figure legends. For statistical test of scRNA-seq dataset see corresponding sections in material and methods.

## ACKNOWLEDGMENTS

We would like to thank A. Brand, B. Edgar, I. Miguel-Aliaga, T. Reiff, P. Patel and J. Veenstra for providing fly lines and reagents. We would like to thank O. Stegle, J. Lohmann, F. Heigwer, A. Coenen-Stass, J. Hawkins and A. Waclawiczek for helpful discussions. We would like to thank the Nikon imaging facility and the Deep sequencing facility at Heidelberg University for support. We thank Florian Heigwer, Alex Henderson, Edward Green and Vaishali Gerwan for providing helpful comments on an earlier version of the manuscript. We thank the Bloomington Drosophila Stock Center and the Vienna Drosophila Resource Center for supplying fly stocks. Computations were run on EMBL IT Services HPC resources.

## AUTHOR CONTRIBUTIONS

Conceptualization: S.R, M.B, W.H. Investigation: S.R., T.W., S.B., L.B., F.P. Single cell RNA-sequencing: S.R., S.L., T.W., S.B., D. I. NanoDam: S.R., S.L., S.P., T.W. Evolutionary analysis: M.H. scRNA-seq data analysis: N.H., E.V., S.P., S.K with input from S.R. Shiny app: N.H. Data analysis: S.R., S.B., W.H., M.B. Supervision: M.B., W.H and S.R. Writing original draft: S.R., M.B., W.H. Writing, reviewing and editing: S.R., N.H., M.B., W.H, with input from all authors.

## DATA AND MATERIAL AVAILABILITY

The code to reproduce all analyses and generate the figures is accessible on GitHub (https://github.com/nickhir/Chronophage). Additionally, the code for the Shiny app is also available on GitHub (https://github.com/nickhir/IntestiMap). Raw sequencing data (FASTQ files) along with the CellRanger output and metadata for each sample is publicly available in the GEO database (GSE276185).

---

**Extended Data Fig. 1.**
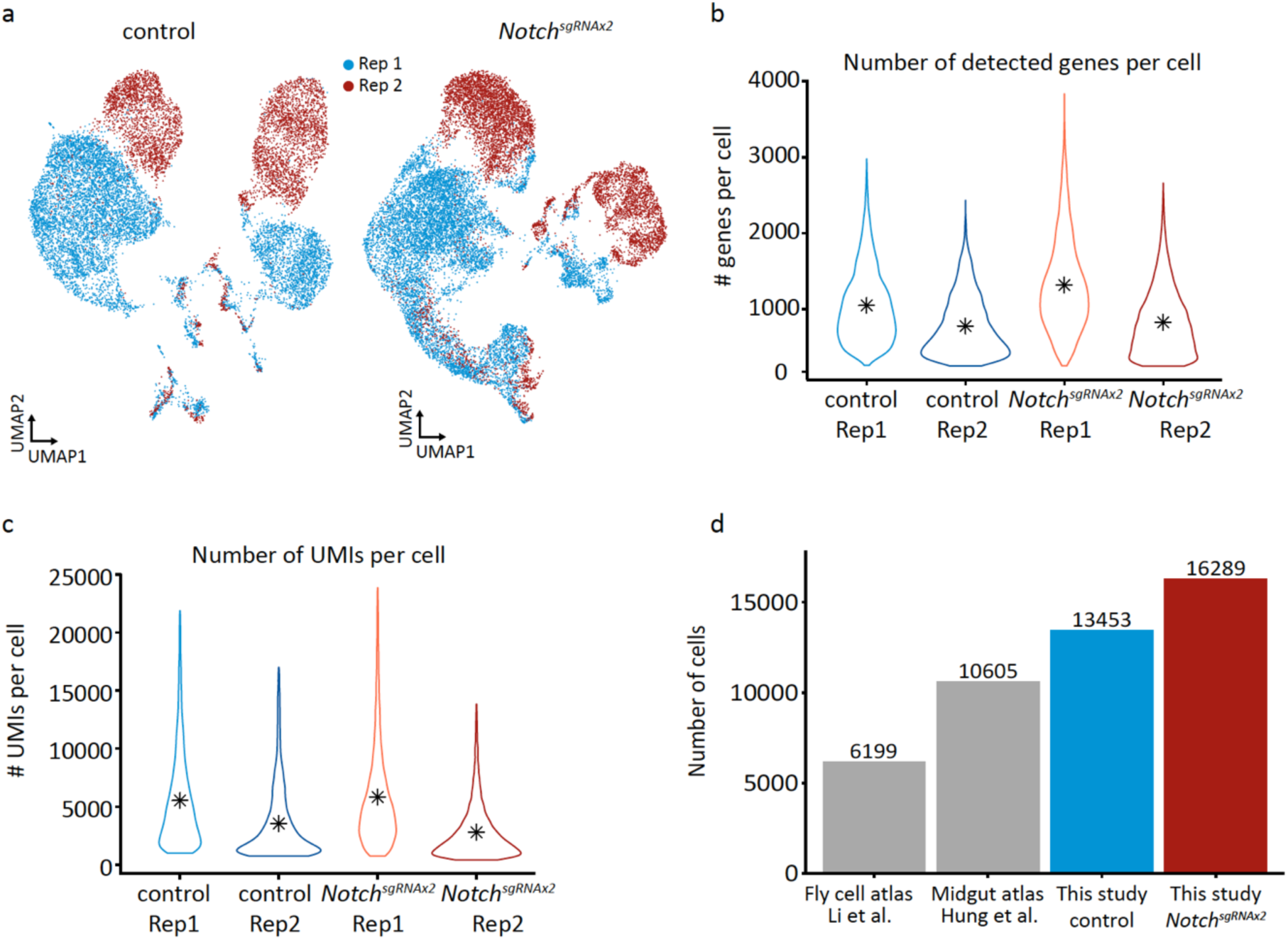
Quality control of scRNA-seq dataset. **(a)** UMAP uncorrected for batch effects for each scRNA-seq replicate. **(b)** Quantification of the number of genes detected per cell across each condition and replicate. **(c)** Quantification of the number of UMIs per cell across each condition and replicate. **(d)** Number of cells recovered after QC from this study compared to two other studies.

**Extended Data Fig. 2.**
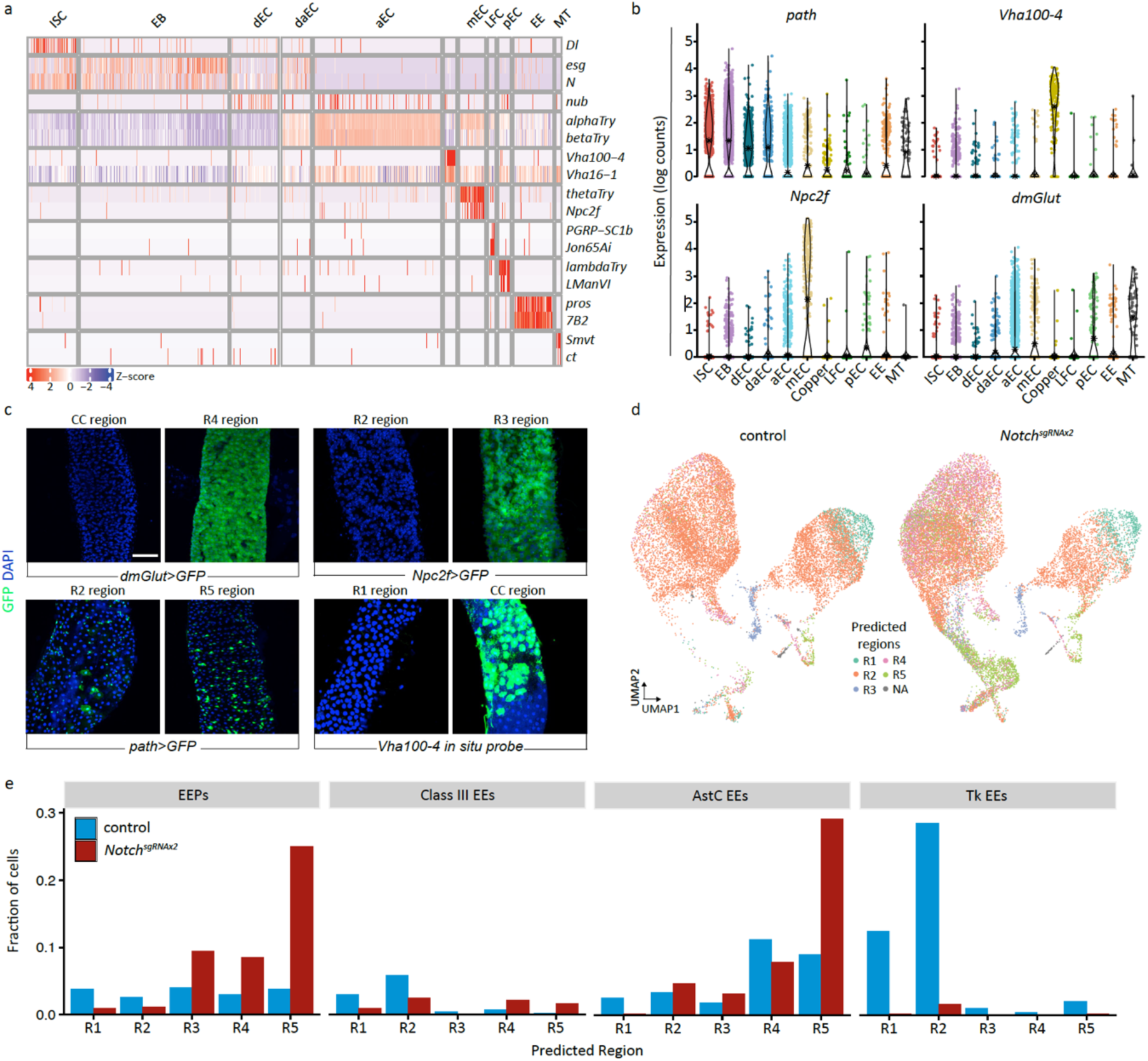
Marker gene expression, regional mapping and velocity profiles of scRNA-seq dataset. **(a)** Z-score of marker gene expression in different intestinal cell types. To preserve space, only a random subset of ISCs, EBs, aECs and dECs was plotted. **(b)** Violin plot of genes with interesting expression profile in different intestinal cell types. **(c)** *in vivo* validation of marker genes in different intestinal regions. **(d)** UMAP of regional prediction of cell types from control and *Notch* mutant condition. **(e)** Regional predictions for EEs and their sub-classes in control and *Notch* mutant condition. Scale bar for c is 100 μm.

**Extended Data Fig. 3.**
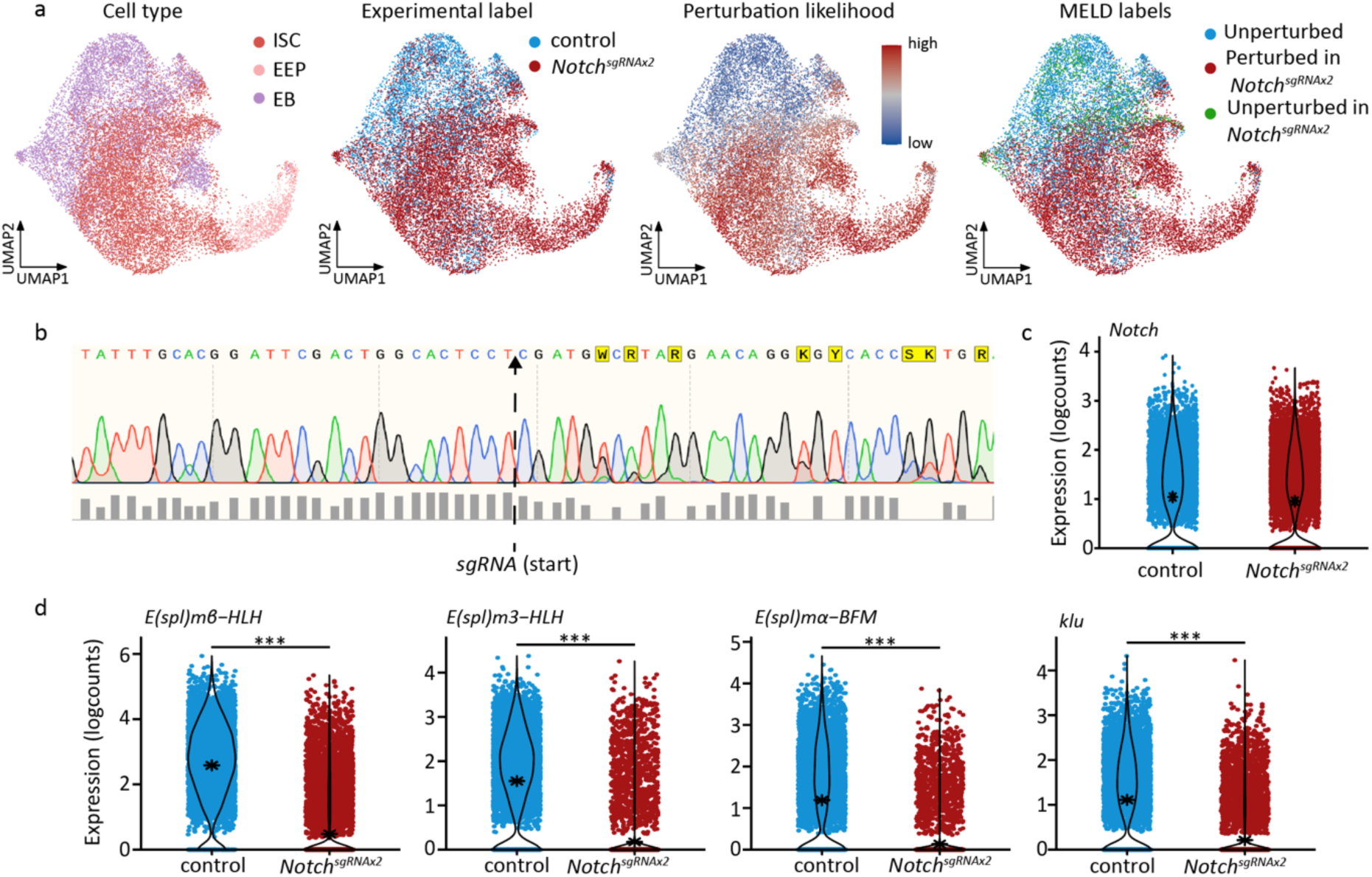
Identification of perturbed cells using MELD. **(a)** MELD UMAPs arranged by cell type, experimental labels, perturbation likelihood were used to generate MELD labels. **(b)** Sanger sequencing traces of progenitor-specific *Notch* mutant intestine, highlighting where *sgRNA* starts. **(c)** Expression of *Notch* in control and Notch mutant condition. Notice the drop in quality (bottom row) after *sgRNA* binding site. **(d)** Expression of various *Notch* target genes in control and *Notch* mutant condition. For all comparisons, an asterisk denotes the mean.

**Extended Data Fig. 4.**
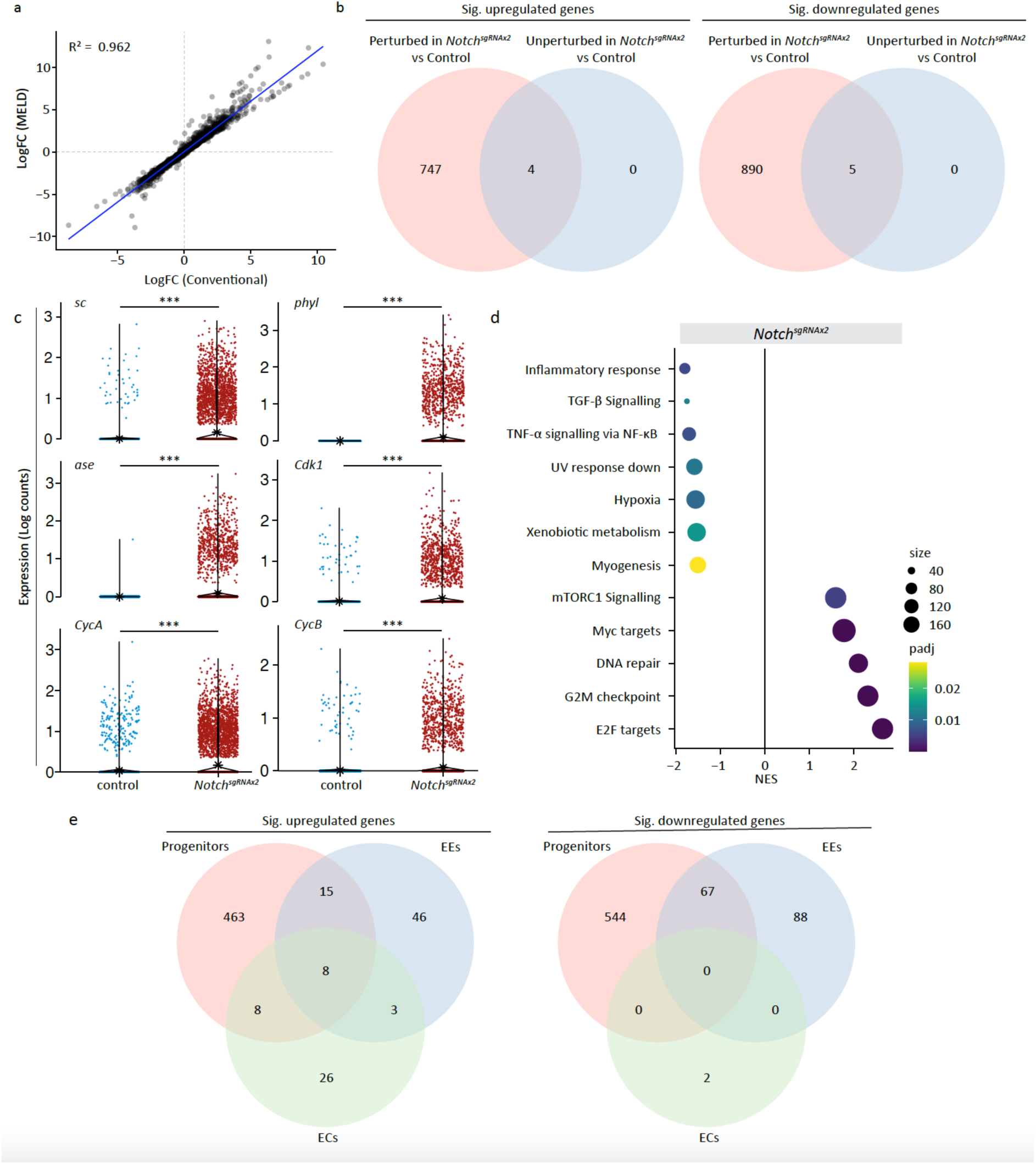
Identification of DEGs based on MELD. **(a)** Correlation coefficient of DEGs identified by MELD and conventional method (see M&M). **(b)** Number of DEGs up and downregulated in perturbed and unperturbed progenitor cells compared to control condition. **(c)** Expression of various genes in control and perturbed *Notch* mutant progenitor cells. **(d)** Hallmark gene set enrichment analysis. **(e)** Number of DEGs that are up and downregulated in different intestinal cell types.

**Extended Data Fig. 5.**
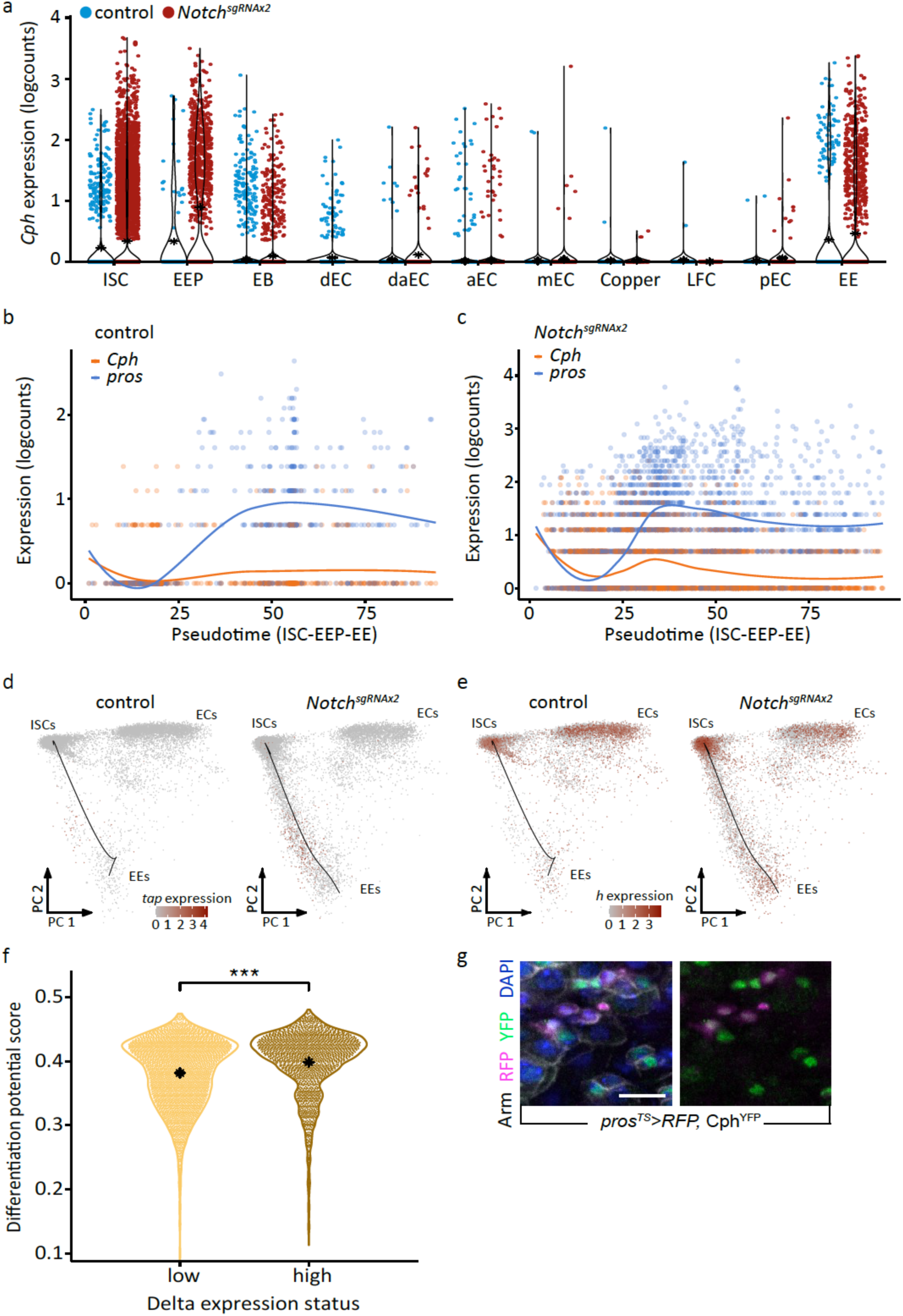
*Cph* expression across intestinal cell types and lineages. **(a)** Expression of *Cph* in all intestinal cell types during homeostasis or *Notch* mutant condition. **(b)** Expression of *Cph* and *pros* during ISC-EEP-EE differentiation in homeostatic condition. **(c)** Expression of *Cph* and *pros* during ISC-EEP-EE differentiation in *Notch* mutant condition. **(d)** PC plot of the expression of *tap* in control and *Notch* mutant condition. **(e)** PC plot of the expression of *h* in control and *Notch* mutant condition. **(f)** Differentiation potential of Dl^High^ and Dl^Low^ ISCs. **(g)** *RFP* expression using the *Pros^TS^* driver overlaps with endogenously tagged Cph^YFP^. Scale bar for g is 10 μm

**Extended Data Fig. 6.**
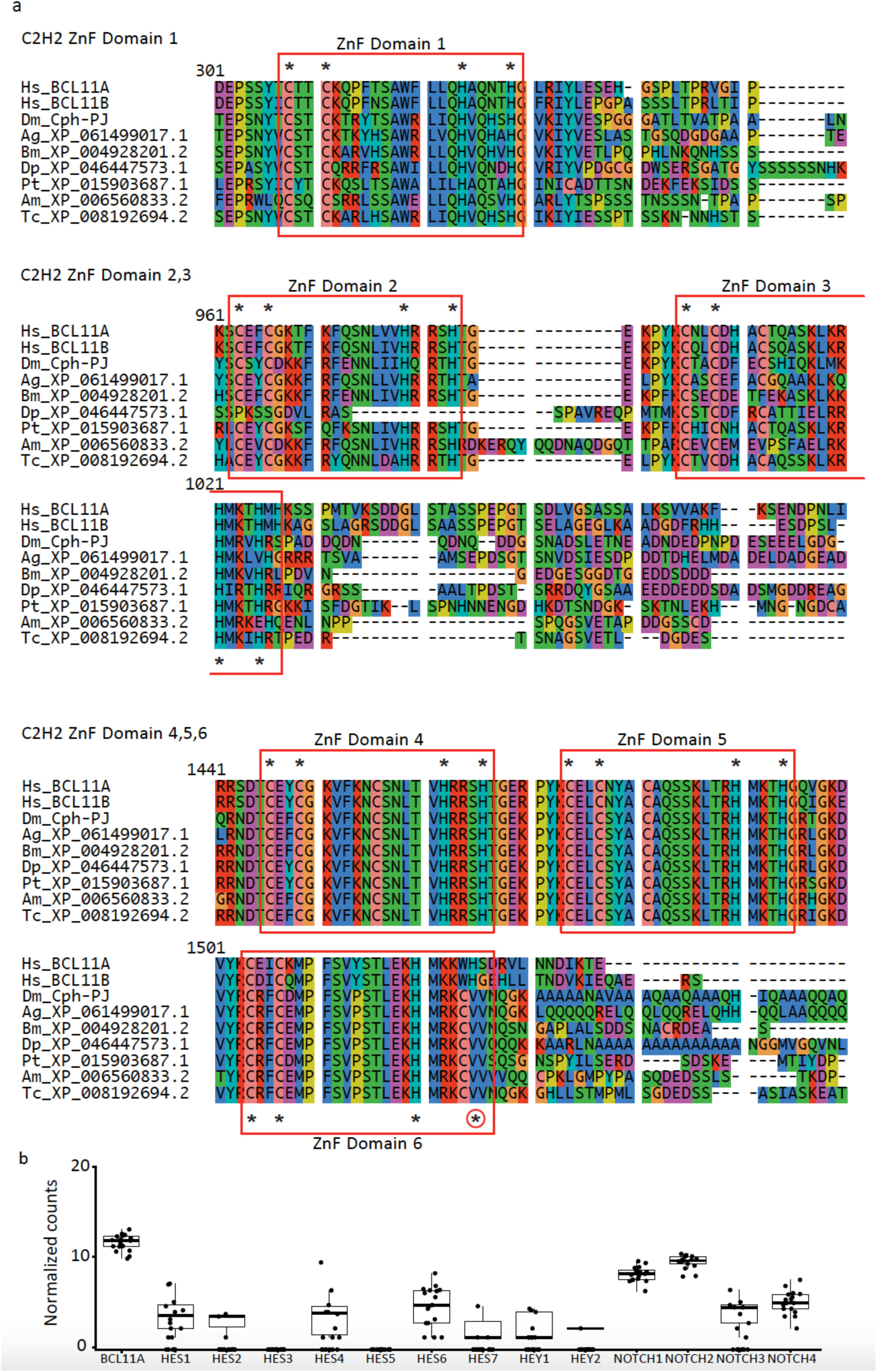
Evolution of BCL11a/Cph and expression in AML. **(a)** Alignment of *Cph* in *Drosophila melanogaster* (Dm) with its homologues in Homo sapiens (Hs), Anopheles gambiae (Ag), Bombyx mori (Bm), Daphnia pulex (Dp), Parasteatoda tepidariorum (Pt), Apis mellifera (Am), and Tribolium castanaeum (Tc). The zinc finger domain (ZnF) of Cph is highly conserved across diverse species. Asterisks denote conserved cysteine or histidine residues, while asterisk inside a circle indicate a lack of conservation across species **(b)** Expression of BCL11a and Notch receptor and downstream target genes in primitive leukemic blasts. Data obtained from patients with AML^39^.

**Extended Data Fig. 7.**
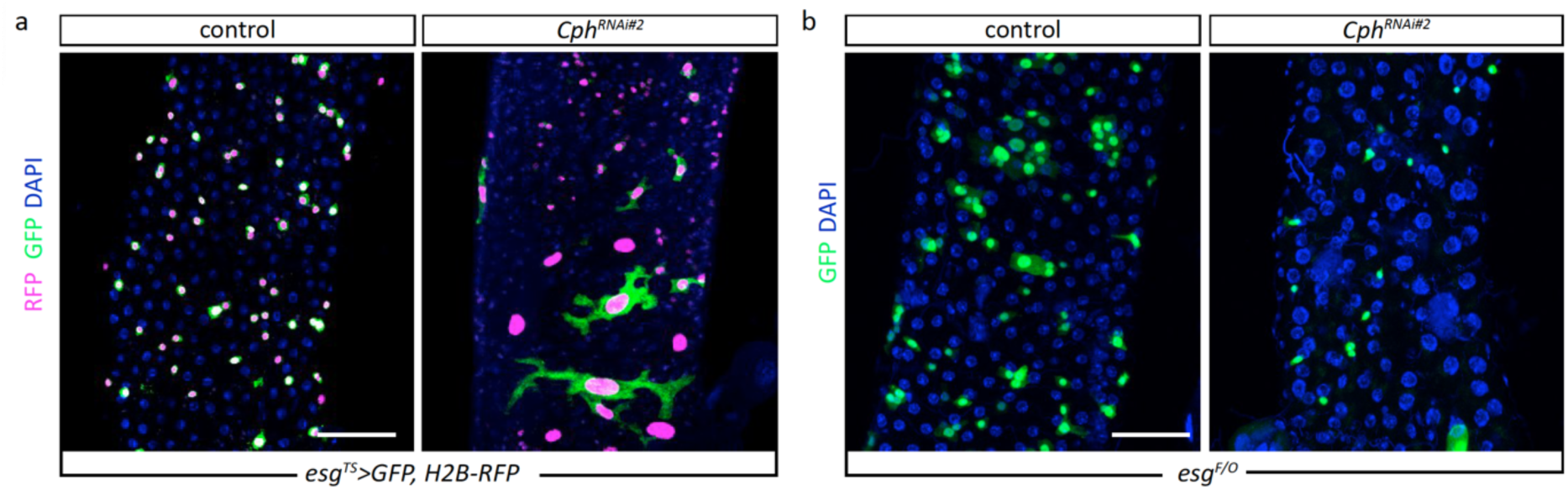
Lineage tracing of Cph depleted progenitor cells. **(a)** REDDM lineage tracing highlighted that *Cph* depletion increases the number *RFP^+^* cells and decreases *GFP^+^/RFP^+^*cell, some of which are abnormally large. Scale bar for all is 100 μm.

**Extended Data Fig. 8.**
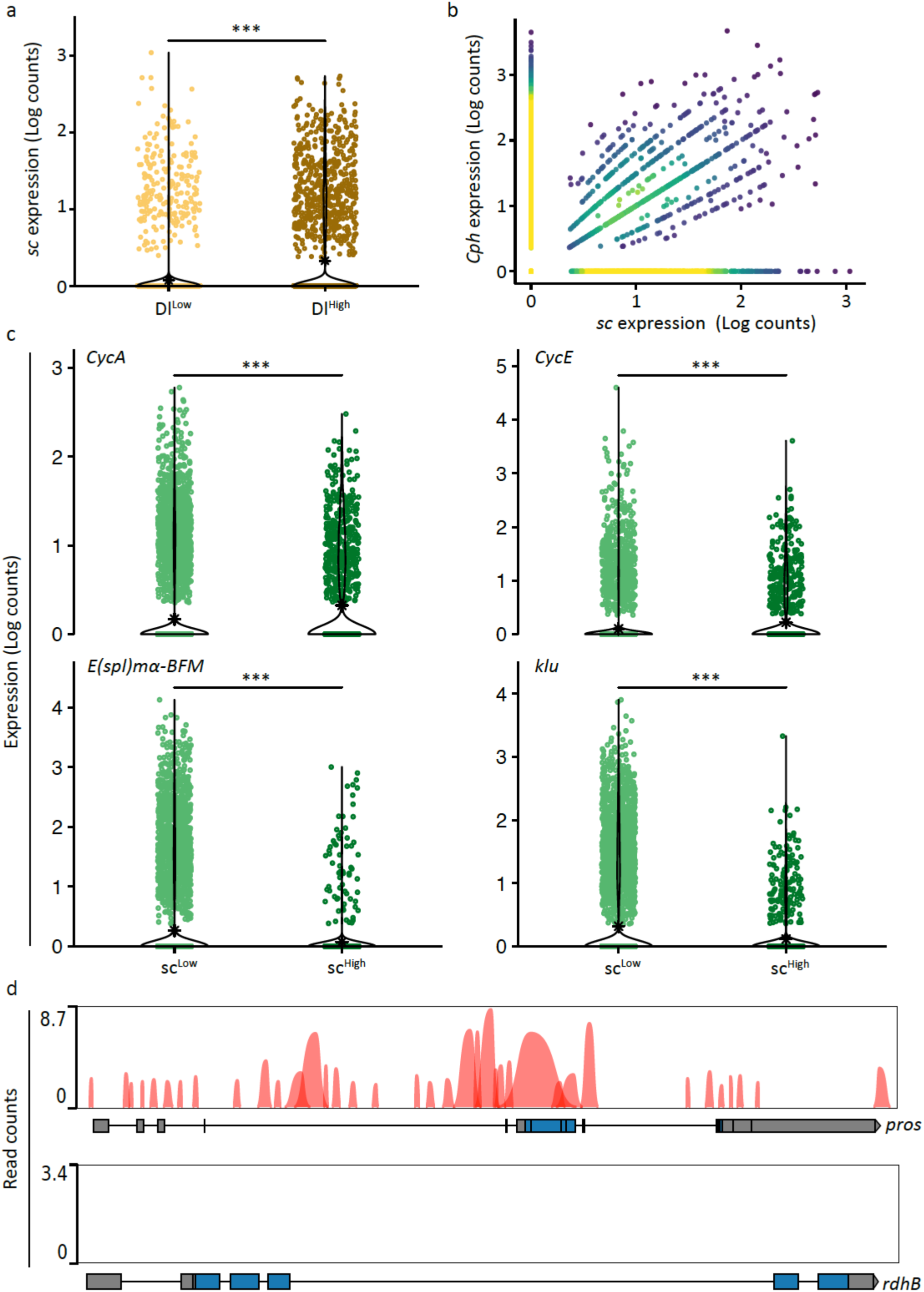
Characteristics of *sc* expressing intestinal cells. **(a)** Expression of *sc* in Dl^High^ and Dl^Low^ ISCs, demonstrating that *sc* is highly expressed in Dl^High^ ISCs. **(b)** Co-expression of *sc* and *Cph*, illustrating high correlation. **(c)** Expression of *CycA*, *CycE*, *E(spl)mα-BFM* and *klu* in sc^High^ and sc^Low^ expressing intestinal cells. **(d)** ChIP-seq tracks for *sc* binding to *pros* and *rdhB*.

**Extended Data Fig. 9.**
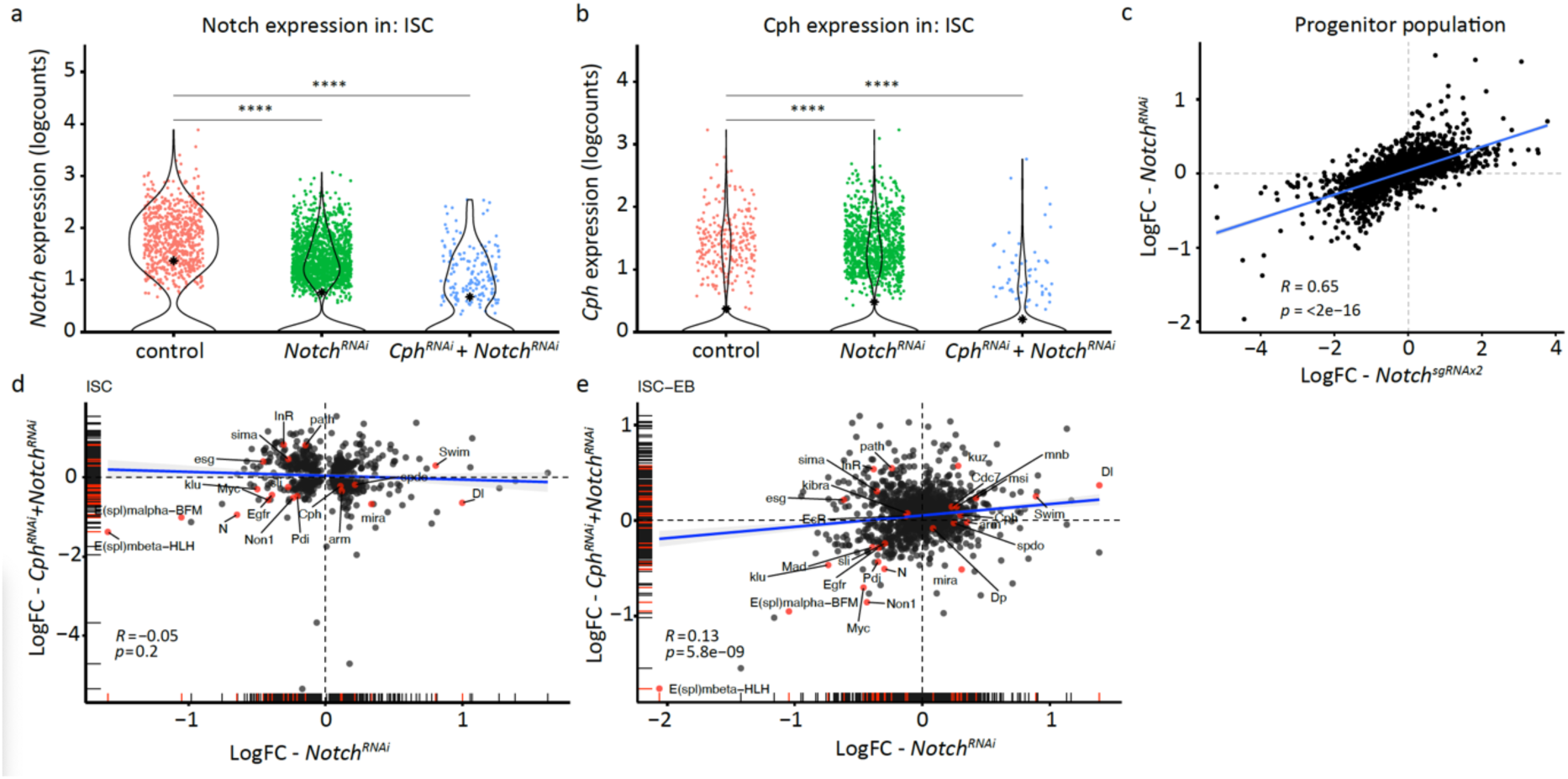
Transcriptional reprogramming of progenitor cells. **(a)** Expression of Notch in ISCs under control, *Notch^RNAi^* and *Cph^RNAi^+Notch^RNAi^* condition. **(b)** Expression of Cph in ISCs under control, *Notch^RNAi^*and *Cph^RNAi^+Notch^RNAi^* condition. **(c)** Correlation coefficient of all DEGs between *Notch^RNAi^* and *Notch^sgRNAx2^*. **(d)** Correlation coefficient of all significant commonly deregulated genes in ISCs in *Notch^RNAi^* and *Cph^RNAi^+Notch^RNAi^*condition. **(e)** Correlation coefficient of all commonly deregulated genes in ISC-EBs in *Notch^RNAi^* and *Cph^RNAi^+Notch^RNAi^*condition.

**Extended Data Fig. 10.**
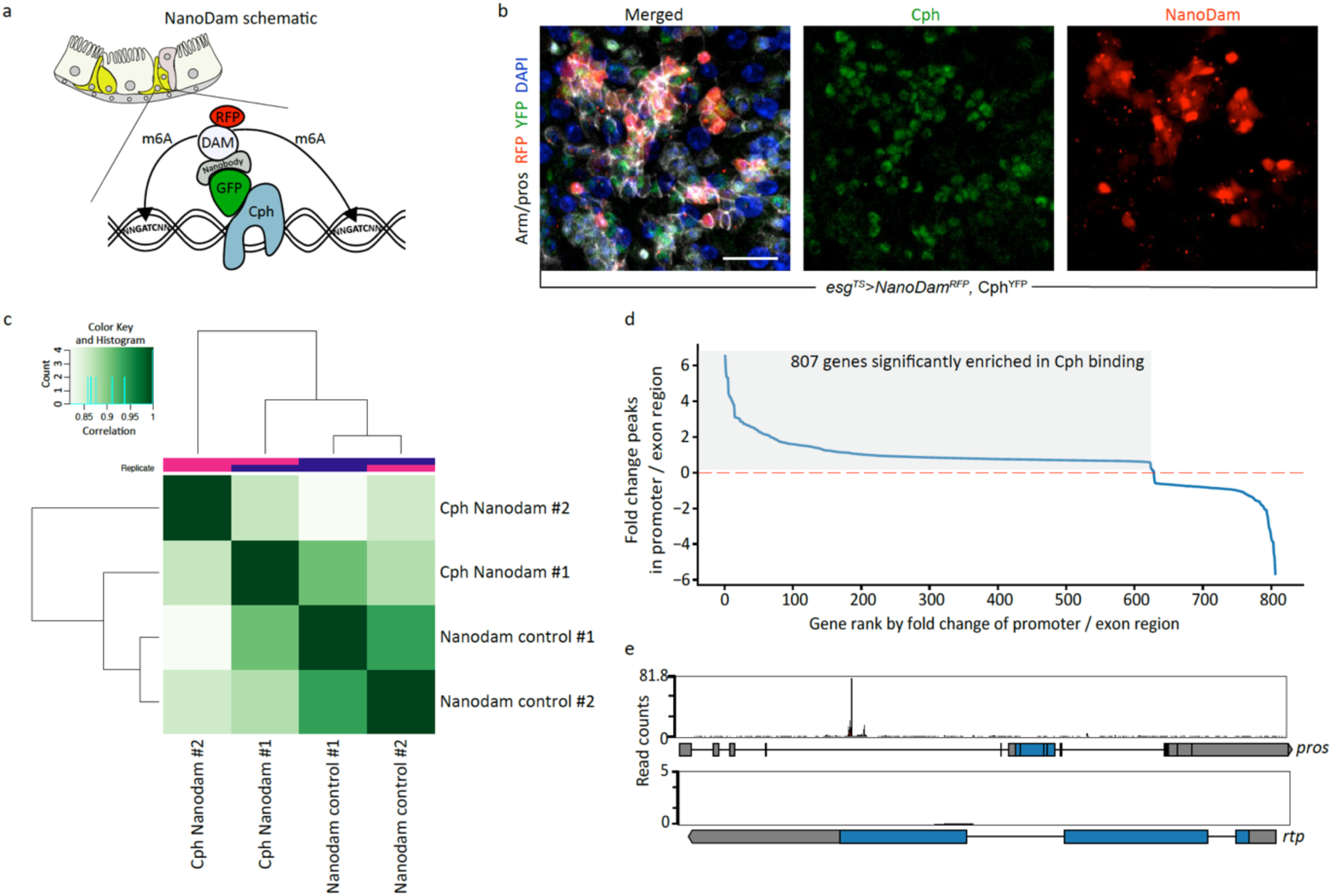
NanoDam profiling of Cph. **(a)** Schematic of NanoDam in progenitor cells. **(b)** Expression of *NanoDam^RFP^* in *GFP^+^* progenitor cells using the *esg^TS^* driver. **(c)** Differential binding correlation heatmap across different replicates. **(d)** Significant Cph target genes within the progenitor population. **(e)** Cph NanoDam binding intensity on the *pros* and *rtp* locus. Notice one major peak within the intronic region of *pros*. Scale bar for b is 100 μm.

